# The propensity to sign-track is associated with externalizing behaviour and distinct patterns of reward-related brain activation in youth

**DOI:** 10.1101/2022.01.29.477945

**Authors:** Janna M. Colaizzi, Shelly B. Flagel, Ashley N. Gearhardt, Michelle A. Borowitz, Rayus Kuplicki, Vadim Zotev, Grace Clark, Jennifer Coronado, Talia Abbott, Martin P. Paulus

## Abstract

Externalizing behaviours in childhood often predict impulse control disorders in adulthood; however, the underlying biobehavioural risk factors are incompletely understood. In animals, the propensity to sign-track, or the degree to which incentive motivational value is attributed to reward cues, is associated with externalizing-type behaviours and deficits in executive control. Using a Pavlovian conditioned approach paradigm, we quantified sign-tracking in healthy 9-12-year-olds. We also measured parent-reported externalizing behaviours and anticipatory neural activations to outcome-predicting cues using the monetary incentive delay fMRI task. Sign-tracking was associated with attentional and inhibitory control deficits and the degree of amygdala, but not cortical, activation during reward anticipation. These findings support the hypothesis that youth with a propensity to sign-track are prone to externalizing tendencies, with an over-reliance on subcortical cue-reactive brain systems. This research highlights sign-tracking as a promising experimental approach delineating the behavioural and neural circuitry of individuals at risk for externalizing disorders.

---

Externalizing tendencies are characterized by a wide range of psychological features, namely impulsivity, defiance, and inattention^1^; and have an early onset^2^, high prevalence rates^2^, and are predictive of impulse control disorders in adulthood.^3^ Thus, understanding the underlying mechanisms at play, particularly in childhood and adolescence, is critical to determine trait vulnerability markers early enough to screen and refer at-risk youth for intervention. A key characteristic of externalizing disorders, and more broadly, impulse control disorders, is increased reactivity to cues.^4^ As such, individual differences in cue-reactivity and associative learning can have significant implications for the development of impulse control disorders^5^ and be used to predict behavioural and mechanistic outcomes relevant to such disorders.^6,7^ In animal studies, behavioural endophenotypes have been identified by individual variation in the propensity to sign-track, or the degree to which animals attribute incentive salience to reward-paired cues.^8–10^ These individual differences are not due to variation in the ability to learn an association, but rather a bias that is evident in a particular conditioned response.^8,9^ Specifically, when presented with a discrete (Pavlovian) cue that has repeatedly been paired with reward, some individuals, goal-trackers (GTs), assign predictive value to the cue and directly approach the location of reward delivery upon cue presentation.^8,10,11^ Others, sign-trackers (STs), approach and interact with the cue itself, thereby attributing both predictive and incentive value to the cue. The attribution of incentive value to the cue renders it attractive and desirable with the ability to capture attention, elicit approach, and promote addiction-related tendencies such as drug-seeking.^8,10,12,13^ Thus, the increased incentive value ascribed to a cue by sign-trackers may be one mechanism by which premature or inappropriate action is initiated and may give rise to vulnerability to impulse control disorders.^5,7,14^ Examining individual differences in the propensity to sign- or goal-track could, therefore, provide an experimental and explanatory translational framework to investigate the neural mechanisms for such disorders.

One candidate process for individual differences in incentive salience attribution and parallel psychopathological vulnerabilities could be the relative imbalance between affective cue-driven and cognitive control systems.^5,14^ Dopamine-dominated subcortical structures facilitate reactive and affectively motivated actions such as fear and reward reactivity whereas cholinergic-dependent cortical structures underlie executive functioning and goal-directed behaviours.^15^ In both humans and animals, the balance between these systems is integral in reward processing and adaptive decision-making, ranging from encoding the value of the reward to economizing the optimal behavioural output to obtain that reward.^14,16^ For goal-tracking rats, a top-down acetylcholine-dominant system directs responses to relevant, goal-oriented stimuli and filters out irrelevant cues, whereas striatal dopamine is crucial for encoding incentive motivational value and a bottom-up, cue-driven, system features more prominently for STs.^10,17–22^ In fact, sign-tracking, but not goal-tracking, can be increased in GTs by inhibition of cortical acetylcholine^19^, while blocking dopamine transmission suppresses sign-tracking but not goal-tracking behaviours.^4,17,23^ This imbalance has implications for variability in impulse control, such that, in rodent STs, a stronger degree of sign-tracking is associated with increased attentional control deficits^24^ and this relationship is modulated by prefrontal acetylcholine.^21^ These patterns are consistent with human impulse control disorders given that (1) striatal dopamine plays a crucial role^4,25,26^ and is deficient^27^ during reward processing and goal-directed tasks for adolescents with impulse control disorders, (2) the ability of reward-paired cues to capture attention is notably stronger for adults with substance use disorder,^28^ and (3) trait impulsivity in youth increases one’s risk for problematic drug and alcohol use.^29^ Moreover, adults who exhibit behaviours related to sign-tracking (e.g., Pavlovian instrumental transfer) also show attentional bias to reward-paired cues.^27^ Thus, the increased propensity to attribute incentive motivational value to reward cues and associated neural mechanisms may put sign-trackers at increased risk of attentional deficits, addiction, and impulse control disorders.

Despite the clear implications sign-tracking has for vulnerability to impulse control disorders, the research in humans is still very sparse, inconsistent, and largely focused on adult populations.^30,31, but see 36^ Various methods have been used to study this behaviour in humans^30,32–35^, however, no prior study has directly applied Pavlovian conditioned approach measures to investigate corresponding neural and trait profiles, and very few have used functional neuroimaging^31^ or assessed the tendency to sign-track in early developmental stages.^36^ Given that adolescence is characterized by a relative imbalance of cortical control in the downregulation of subcortical systems^37^, for some, sign-tracking may dominate the adolescent brain. Moreover, the considerable overlap between sign-tracking and externalizing traits suggests that examining these phenotypes in youth, prior to typical symptom onset for impulse control disorders, offers the potential to delineate the neural circuitry that is associated with a vulnerability or risk for externalizing disorders early enough to implement preventative and interventive measures; thus, reinforcing the prospective translational utility of the model. We therefore developed a Pavlovian conditioned approach paradigm relevant to pre-adolescent youth to determine whether sign-tracking and goal-tracking behaviours could be identified in this population. Further, we measured externalizing tendencies and anticipatory neural responses to reward and loss, hypothesizing that those with biased approach behaviours to the CS (STs), would show neural and behavioural profiles consistent with both sign-tracking in animals and externalizing disorders in youth.

## Results

### Pavlovian conditioned approach

Our primary goal was to establish a Pavlovian conditioning paradigm comparable to those used in animal models to identify sign-tracking and goal-tracking phenotypes in youth. We used the Pavlovian conditioning paradigm described by Flagel and colleagues (2008)^38^ for rodents and adapted for humans by Joyner and colleagues (2018)^36^ as a basis for the current study (Figure 1a). Participants (N = 40, Table 1; see Methods for a priori power analysis report) completed 40 response-independent conditioning trials consisting of lever (CS) presentation and subsequent reward (US; $0.20 token; all monetary units in USD) presentation. A randomly selected inter-trial interval (ITI) followed each CS-US trial.

**Figure 1.**
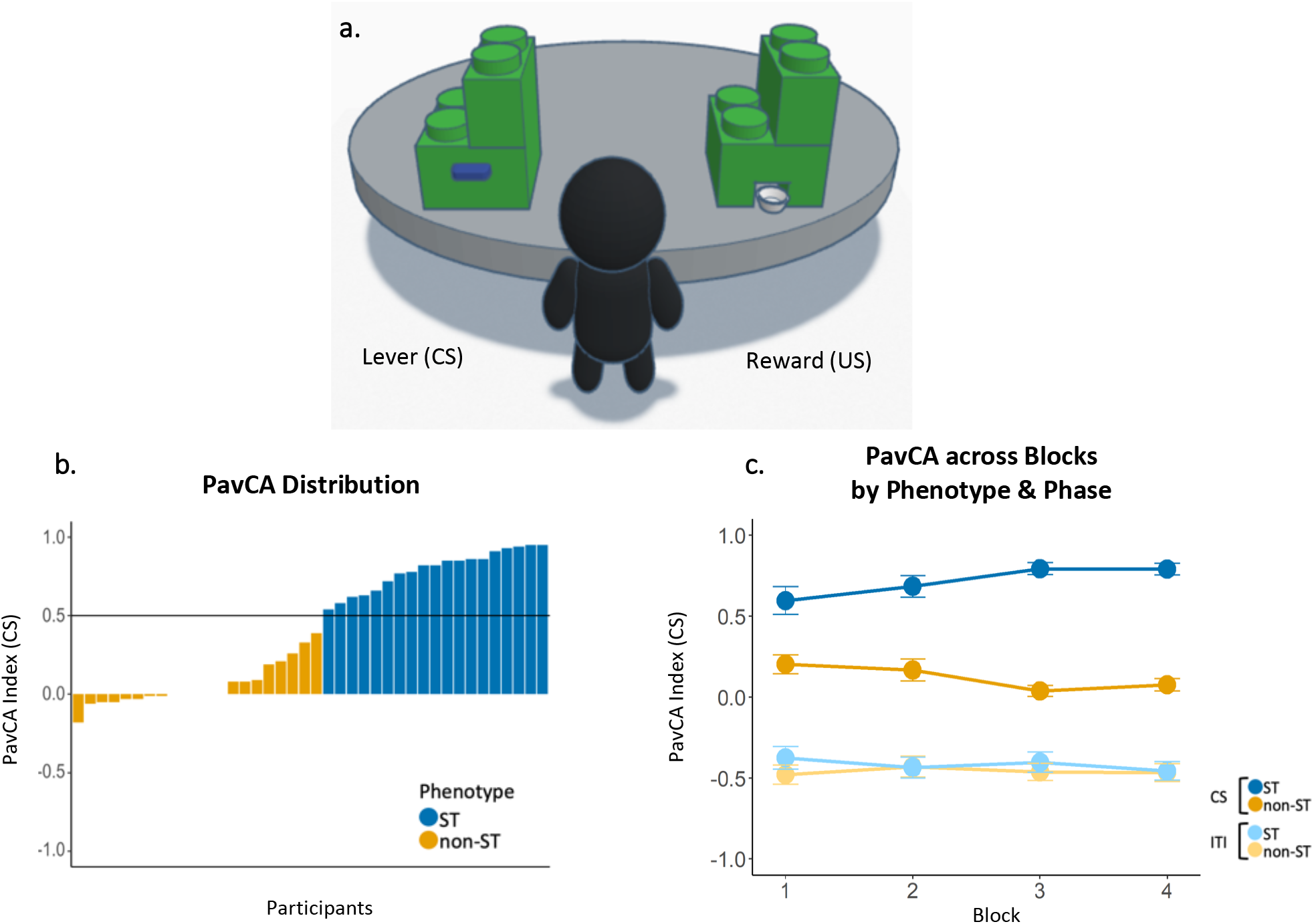
Human Pavlovian conditioned approach apparatus and behaviors. **a.** Digital depiction of the Pavlovian Conditioned Approach task adapted for use in human youth. Lever response box shown on the left and reward response box on the right. CS = conditioned stimulus; US = unconditioned stimulus. **b.** PavCA = Pavlovian Conditioned Approach; Individual subject distribution of PavCA index scores averaged across Blocks 3 and 4. Phenotypes were characterized by a PavCA score greater than 0.5 (horizonal line) for sign-trackers and less than 0.5 for non-sign-trackers; ST = sign-tracker; non-ST = non-sign-tracker. **c.** PavCA scores across all 4 blocks between phenotypes (ST, non-ST) and between phases (CS, ITI); CS = conditioned stimulus/lever presentation; ITI = inter-trial interval. Error bars represent standard error of the mean. Post-hoc tests between phenotypes during CS for each block are marked with asterisks; *p* < .001***, *p* < .01**, *p* < .05*

**Table 1.**
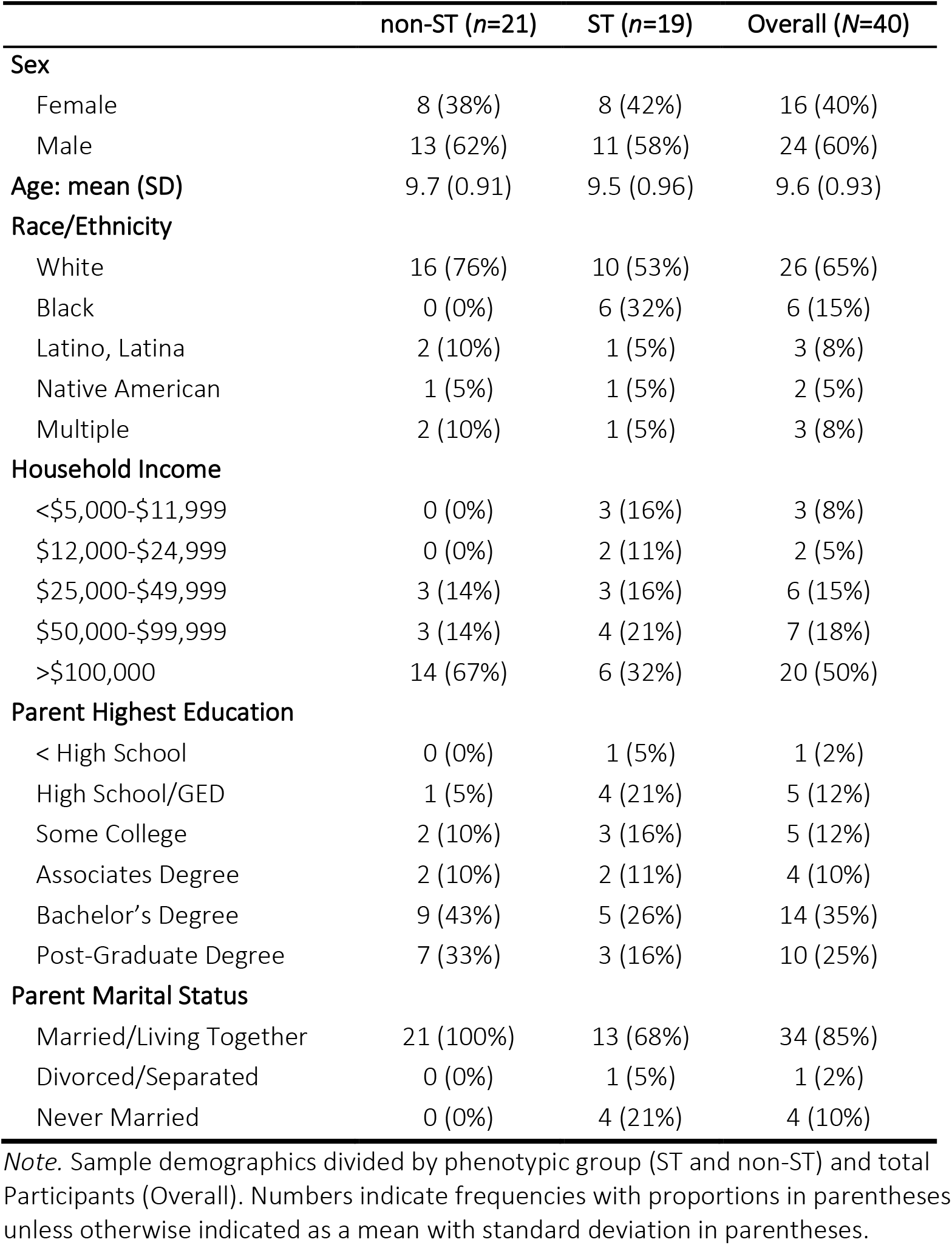
Proportions and means/standard deviations for sample demographics by phenotype.

|  | non-ST (*n*=21) | ST (*n*=19) | Overall (*N*=40) |
| --- | --- | --- | --- |
| **Sex** |  |  |  |
| Female | 8 (38%) | 8 (42%) | 16 (40%) |
| Male | 13 (62%) | 11 (58%) | 24 (60%) |
| **Age: mean (SD)** | 9.7 (0.91) | 9.5 (0.96) | 9.6 (0.93) |
| **Race/Ethnicity** |  |  |  |
| White | 16 (76%) | 10 (53%) | 26 (65%) |
| Black | 0 (0%) | 6 (32%) | 6 (15%) |
| Latino, Latina | 2 (10%) | 1 (5%) | 3 (8%) |
| Native American | 1 (5%) | 1 (5%) | 2 (5%) |
| Multiple | 2 (10%) | 1 (5%) | 3 (8%) |
| **Household Income** |  |  |  |
| <\$5,000-\$11,999 | 0 (0%) | 3 (16%) | 3 (8%) |
| \$12,000-\$24,999 | 0 (0%) | 2 (11%) | 2 (5%) |
| \$25,000-\$49,999 | 3 (14%) | 3 (16%) | 6 (15%) |
| \$50,000-\$99,999 | 3 (14%) | 4 (21%) | 7 (18%) |
| >\$100,000 | 14 (67%) | 6 (32%) | 20 (50%) |
| **Parent Highest Education** |  |  |  |
| < High School | 0 (0%) | 1 (5%) | 1 (2%) |
| High School/GED | 1 (5%) | 4 (21%) | 5 (12%) |
| Some College | 2 (10%) | 3 (16%) | 5 (12%) |
| Associates Degree | 2 (10%) | 2 (11%) | 4 (10%) |
| Bachelor's Degree | 9 (43%) | 5 (26%) | 14 (35%) |
| Post-Graduate Degree | 7 (33%) | 3 (16%) | 10 (25%) |
| **Parent Marital Status** |  |  |  |
| Married/Living Together | 21 (100%) | 13 (68%) | 34 (85%) |
| Divorced/Separated | 0 (0%) | 1 (5%) | 1 (2%) |
| Never Married | 0 (0%) | 4 (21%) | 4 (10%) |

*Note.* Sample demographics divided by phenotypic group (ST and non-ST) and total Participants (Overall). Numbers indicate frequencies with proportions in parentheses unless otherwise indicated as a mean with standard deviation in parentheses.

Feasibility was assessed by identifying both qualitative (Supplemental Figure S1) and quantitative markers of child engagement and learning. We identified individual differences in the attribution of incentive salience to reward-paired cues (sign-tracking or goal-tracking behaviours) via a Pavlovian Conditioned Approach (PavCA) index based on previously used models in animal studies^39^ derived from the number and timing of physical contacts to the CS and US during CS-presentation and ITI phases.

Consistent with animal models^4^, we classified categorical phenotypic groups using a PavCA value during the CS-period of 0.5 or greater (STs) and less than −0.5 (GTs). PavCA scores during the CS-period ranged from −0.18 to 0.95 (*m* = 0.41, *sd* = 0.40; Supplemental Table S1) and the distribution indicated that participant behaviours were skewed toward either neutral or lever-directed behaviours (Figure 1b), therefore we were able to identify STs, but not GTs. Nineteen participants were identified as STs (*mp_PavCA_* = 0.79, *sd* = 0.13), and since no participants had a PavCA value less than −0.5, we classified the remaining 21 participants as non-sign-trackers (non-ST; *m_PavCA_* = 0.06, *sd* = 0.14) and used these as our comparison groups (see Supplemental Materials for video examples of ST and non-ST behaviours). Scores did not differ by sex (*t*_37.8_ = −0.48, *p* = .633, *d* = 0.15, s.e. = 0.13, 95% CI = −0.32–0.20) or age (*t*_37.8_ = 0.22, *p* = .824, *d* = 0.07, s.e. = 3.91, 95% CI = −7.05–8.80).

To assess learning of conditioned responses to the CS, we examined behavioural responses to the CS using two-sided linear mixed effects models with the factors time (Block 1-4), phase (CS, ITI), and phenotype (ST, non-ST) for lever- and reward-directed behaviours (Supplemental Figure S2). Because CS lever contacts correlated positively with ITI lever contacts (*r* = 0.59, *p* < .001, Supplemental Table S2), a normalised frequency score was calculated for lever- and reward-directed behaviours (contacts and probability) in order to adequately compare between CS and ITI periods. Briefly, we observed that STs demonstrated a higher probability to contact the lever during the CS period and over time (three-way interaction, *F*_3,266_ = 3.84, *p* = .010, *η*^2^_p_ = 0.04, 90% CI = 0.00 – −0.08). STs and non-STs differed in their probability to contact the reward between phases (phase main effect, *F*_1,266_ = 504.79, *p* < .001, *η*^2^_p_ = 0.65, 90% CI = 0.60–0.70) however there were no significant phenotypic differences during either phase or over time (three-way interaction, *F*_3,266_ = 2.07, *p* = .104, *η*^2^_p_ = 0.02, 90% CI = 0.00–0.05). Of note, there were significant main effects for age (*F*_1,36_ = 6.62, *p* = .010, *η*^2^_p_ = 0.16, 90% CI = 0.02–0.34) and sex (females higher; *F*_1,36_ = 7.43, *p* = .009, *η*^2^_p_ = 0.17, 90% CI = 0.03–0.35) in the probability to contact the reward which may implicate developmental differences impacting goal-tracking behaviours. Together, it appears that much of the individual variation in behaviour stemmed from lever-directed behaviours during CS presentation which supports the general skew towards sign-tracking behaviour.

We also examined PavCA behaviour between phases for each phenotype over time using non-normalised metrics (Figure 1c). PavCA scores showed significant main effects for phenotype (*F*_1,36_ = 52.82, *p* < .001, *η*^2^_p_ = 0.59, 90% CI = 0.42–0.71) and phase (*F*_1,265.1_ = 1274.34, *p* < .001, *η*^2^_p_ = 0.83, 90% CI = 0.80– 0.85), suggesting that participants are behaviourally discriminating between phases, and STs are doing so to a greater degree. STs increasingly approached the lever over time (higher PavCA score; phenotype by time interaction, *F*_3,265.1_ = 2.63, *p* = .050, *η*^2^_p_ = 0.03, 90% CI = 0.00–0.06) and more so during CS presentation (phenotype by phase interaction, *F*_1,265.1_ = 131.46, *p* < .001; three-way interaction; *F*_1,265.1_ = 4.21, *p* = .006, *η*^2^_p_ = 0.05, 90% CI = 0.01–0.09; Figure 2c). This appears to reflect learning of the conditioned response and the characterisation of sign-tracking by selectively increasing lever approach during the CS period. Conversely, non-STs progressively decrease their lever-directed behaviours over the course of training, further characterising the distinction between phenotypic responses. PavCA showed a significant main effect for sex (males higher; *F*_1,36.1_ = 13.28, *p* = .001, *η*^2^_p_ = 0.27, 90% CI = 0.09–0.45) but not age (*F*_1,36_ = 1.64, *p* = .208, *η*^2^_p_ = 0.04, 90% CI = 0.00–0.19). Together, these results show that the two phenotypes selectively discriminate between phases and, during CS, PavCA behaviours continually diverge over the course of the session.

**Figure 2.**
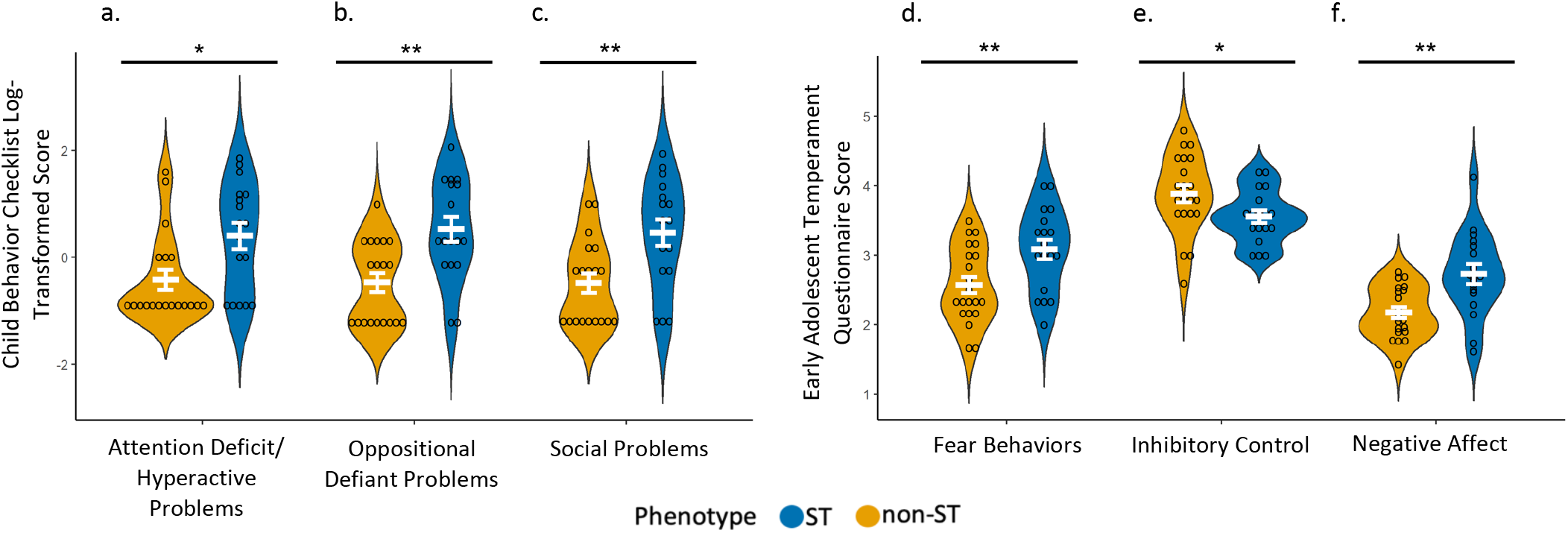
Violin plots illustrating phenotypic differences in externalizing behaviors. Plotted results of two-sided Welch two-sample *t*-tests measuring mean differences between phenotypes on parent-reported externalizing behaviors. ST = sign-trackers (n = 19); non-ST = non-sign-trackers (n = 21). *p* < .01**, *p* < .05* **a.-c.** Scores from the Child Behavior Checklist (CBCL) subscales log transformed. **a.** Scores from the CBCL DSM-oriented subscale for attention deficit/hyperactive problems. **b.** Scores from the CBCL DSM-oriented subscale for oppositional defiance. **c.** Scores from the CBCL subscale for social problems includes items relating to jealousy, not getting along with others, and not being liked by others. **d.-f.** Items on the Early Adolescent Temperament Questionnaire (EATQ). **d.** Scores from the EATQ fear subscale address unpleasant affect related to anticipation of distress. **e.** Scores on the inhibitory control subscale of the EATQ address the capacity to plan and suppress inappropriate responses. **f.** Negative affect scores on the EATQ is a composite score of aggression, depression, and frustration subscales.

### Externalizing behaviours by phenotype

The data presented so far provides evidence for a bias toward responding to reward-paired cues, indicative of sign-tracking, in a subset of youth. Given the well-established characteristic differences between STs and GTs in animal studies^14,21^, we aimed to validate the application of phenotypic distinctions in human youth. Our primary hypothesis was that human STs would demonstrate symptomatic and neurobiological profiles consistent with externalizing characteristics and a reliance on bottom-up processing rather than top-down cognitive control. We examined this using developmentally validated parent-report questionnaires (Child Behaviour Checklist, CBCL^40^ and Early Adolescent Temperament Questionnaire, EATQ^41^). It is important to note that these measures do not represent clinical diagnoses, but dimensions of behaviour associated with psychiatric symptoms. Measures were tested for normality and all CBCL subscales were log transformed. To directly compare measures between STs and non-STs, we used two-sided Welch two sample *t*-tests (Supplemental Table S3; Figure 2) with FDR *p*-value corrections used for multiple comparisons.

Given that attentional control deficits are a hallmark characteristic in rodent STs,^19,21^ and given the shared circuitry between sign-tracking rats^21^ and humans with attention-based and impulse control disorders,^42^ we expected elevated attention problems in the sign-trackers in our sample. Indeed, when compared to non-STs, STs showed increased symptoms of attention deficit/hyperactivity problems (CBCL; *t*_30.9_ = −2.66, *p* = .014, *d* = 0.90, s.e. = 0.31, 95% CI = −1.45 – −0.19; Figure 2a) which supports prior work highlighting differences in reward processing in attention disorders.^43^

Animal literature has further identified characteristic differences in fear responses, such that, when exposed to fear conditioning tasks, STs show exaggerated fear-associated cue reactivity consistent with an increased susceptibility to both substance use and post-traumatic stress.^44,45^ When compared to non-STs, STs in our sample had increased reported symptoms of fear (EATQ; unpleasant affect related to anticipation of distress; *t*_35_ = −2.79, *p* = .012, *d* = 0.90, s.e. = 0.18, 95% CI = −0.88 – −0.14; Figure 2d), which is in line with previous animal findings implicating STs as having an increased vulnerability to fear-related responses to cues, regardless of context.^45^

Furthermore, inhibitory control deficits have been associated with sign-tracking in rats^18,20,46^ as well as an increased vulnerability for impulse control disorders and addiction in humans.^47^ In our sample, STs had lower scores on indices of inhibitory control (EATQ; the capacity to plan and suppress inappropriate responses; *t*_35.2_ = 2.17, *p* = .037, *d* = 0.68, s.e. = 0.15, 95% CI = 0.02–0.64; Figure 2e); however, the degree of inhibitory control was not significantly correlated with PavCA scores (*r* = −0.18, *p* = .290; Supplemental Table S4) and there were no group differences in self-reported measures of behavioural inhibition/impulsivity or behavioural measures of impulsivity (see Methods section for details; Supplemental Table S3). Given previous findings^16,18,48^, this relationship should be further examined within a larger sample.

Beyond the translational utility of capturing sign-tracking tendencies in youth, we aimed to examine symptomatic differences between phenotypes consistent with human-specific psychopathology. Specifically, we expected to see behavioural tendencies in STs that are developmentally characteristic of externalizing disorders. In contrast to non-STs, caregivers reported STs as having increased oppositional defiant problems (CBCL; *t*_31_ = −3.44, *p* = .006, *d* = 1.16. s.e. = 0.29, 95% CI = −1.58 – −0.40; Figure 2b) and social problems (CBCL; including items relating to jealousy, not getting along with others, and not being liked by others; *t*_30.2_ = −3.10, *p* = .008, *d* = 1.05, s.e. = 0.30, 95% CI = −1.56 – −0.32; Figure 2c). Although these two subscales are positively correlated (*r* = 0.64, *p* < .001; Supplemental Table S4) suggesting homogenous traits, they are derived from independent items on the CBCL. STs also showed increased levels of negative affect (EATQ composite score of frustration, depressive mood, and aggressive behaviours, *t*_27.3_ = −3.14, *p* = .006, *d* = 1.14, s.e. = 0.16, 95% CI = −1.89 – −0.22; Figure 2f). Taken together, the association between the degree of sign-tracking and these measures support the notion that STs exhibit deficits in behavioural regulation and inhibitory control as well as a bias toward affective/reactive responding across multiple diagnostic criteria.^48^

### Neuroimaging

The results presented above demonstrate behaviours and tendencies specific to STs that may broadly indicate a reliance on bottom-up processing and are consistent with characteristics of both rodent models and theoretical accounts of translation of these paradigms to clinical populations.^49^ To further elucidate the neurobiological processes that may contribute to the propensity to sign-track, we employed functional neuroimaging to measure reward processing. Participants completed the Monetary Incentive Delay (MID) fMRI task^50^ to determine the brain response to a gain, no gain, or loss predictive cue. We performed a whole-brain voxelwise linear mixed effects analysis with fixed effects for group, condition, age, sex, a group by condition interaction and a random intercept for subject, followed by planned contrasts investigating a group (ST, non-ST) by condition (win vs neutral or loss vs neutral) interaction. Estimated marginal means were used for post hoc tests with Tukey *p*-value adjustment for multiple comparisons. A total of 29 subjects (n_ST_ = 12, n_non-ST_ = 17) survived motion correction.

When examining the win-neutral contrasts during the MID anticipatory phase, BOLD activations in the left inferior parietal lobe (IPL) showed a significant group by condition interaction (*F*_1,27_ = 10.30, *p* = .003, *η*^2^_p_ = 0.28, 90% CI = 0.07–0.48, Figure 3a; Supplemental Table S5 for details on coordinates and volumes for these and additional significant regions). Post hoc tests indicate that non-STs significantly increase activation from neutral to large win conditions (*t*_27_ = −4.15, *p* = .002, *d* = −1.42, s.e. = 0.37, 95% CI = −2.19 – −0.66), indicating salience-dependent modulation within the IPL. Conversely, there was no evidence for a similar effect in STs and activations did not differ between conditions or from non-STs during the win condition (*p*s > 0.8). Importantly, this same modulatory pattern is evident in the left IPL for the loss-neutral contrast during the anticipatory phase (group by condition interaction; *F*_1,27_ = 12.31, *p* = .002, *η*^2^_p_ = 0.31, 90% CI = 0.09–0.51; Figure 3b), which appears to indicate that this effect is not reward-specific but rather salience-driven. This same pattern is consistent across multiple cortical regions implicated in cognitive control (Supplemental Table S5) which may indicate that, in comparison to non-STs, STs do not actively engage cognitive control and the salience of the cue is cognitively irrelevant.

**Figure 3.**
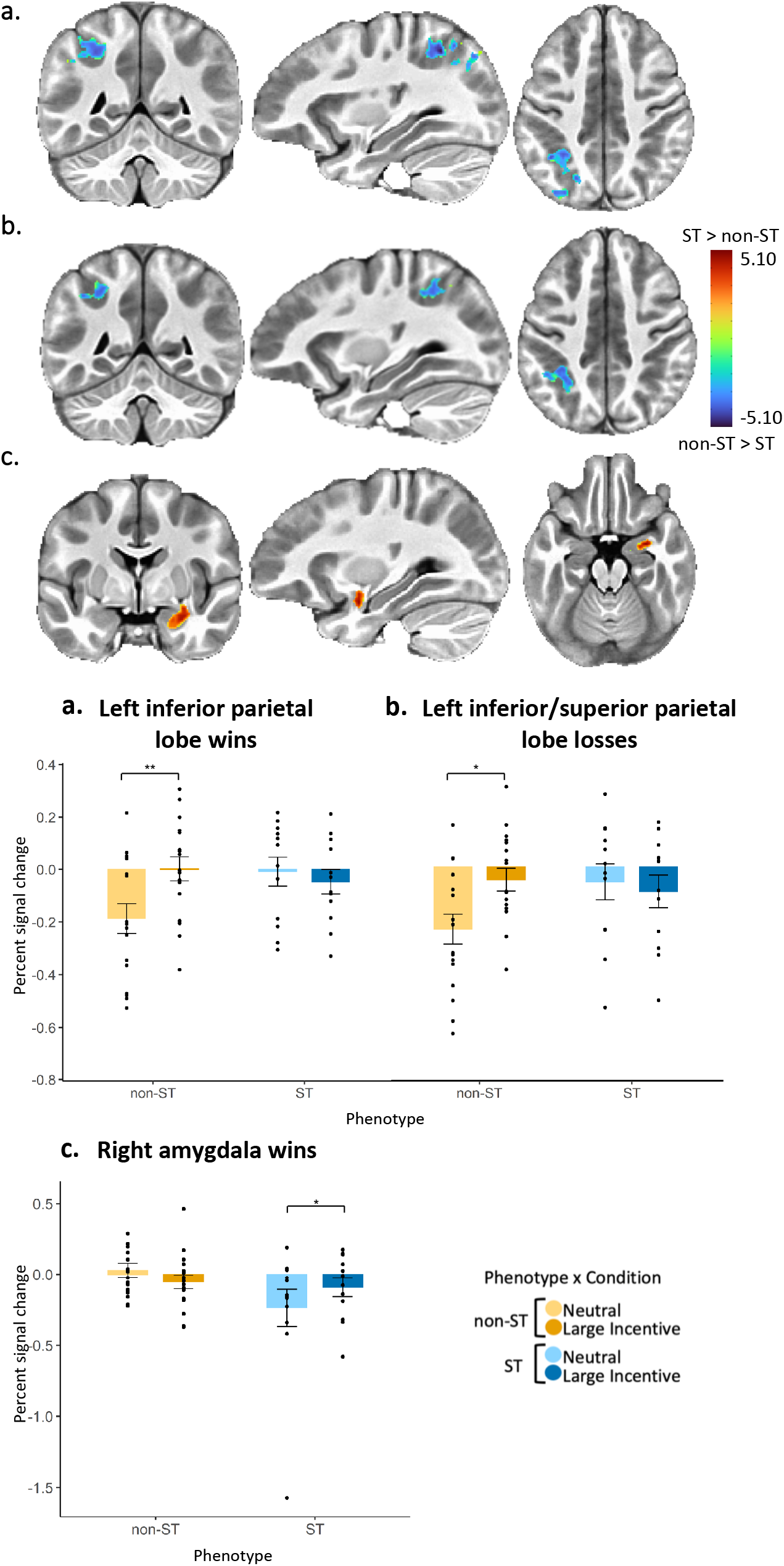
Whole brain voxel-wise functional magnetic resonance imaging (fMRI) percent BOLD signal change during the Monetary Incentive Delay (MID) task for win and loss anticipation. ST = sign-tracker (n = 12), non-ST = non-sign-tracker (n = 17). Participants were presented with an anticipatory cue indicating the upcoming potential gain, no gain, or loss. Results are based on whole-brain voxelwise linear mixed effects with fixed effects for group, condition, age, sex, a group by condition interaction and a random intercept for subject. We followed this with planned contrasts investigating a group (ST, non-ST) by condition (win-neutral or loss-neutral) interaction. Significant clusters are reported using a voxelwise *p*-value threshold of 0.005 and α < 0.05 at the cluster level (N = 11.73 voxels). Estimated marginal means were used for post-hoc tests. Brain activation colors represent *t*-test *z*-statistics for group differences in percent signal change between respective conditions (win-neutral, loss-neutral). Blue represents negative values and indicates a greater percent signal change for non-ST than ST between neutral and large incentive conditions (win/loss). Red represents positive values that indicate greater percent signal change for STs between neutral and large incentive conditions. Bar graphs depict percent signal change in respective clusters across neutral and large incentive conditions. Non-STs are represented by yellow bars and STs by blue. Lighter colors indicate neutral trials whereas darker colors indicate win/loss trials. Significant differences derived from post hoc tests are marked with asterisks; *p* < .01**, *p* < .05*. Significant contrasts (win-neutral or loss-neutral) correspond with brain images such that blue indicates a significant contrast for non-ST and red indicates a significant contrast for STs. **a.** Blood oxygen level dependent (BOLD) activations for a group by condition interaction in the left inferior parietal lobe during win anticipation compared to neutral anticipation. Post hoc tests indicate that non-STs significantly increase BOLD activation from neutral to large win conditions but STs do not. STs marginally differed from non-STs during neutral but not during high win. **b.** BOLD activations for a group by condition interaction in the left inferior parietal lobe during loss anticipation compared to neutral anticipation. Post hoc tests indicate that non-STs significantly increase BOLD activation from neutral to loss conditions but STs do not. **c.** BOLD activations for a group by condition interaction in the right amygdala during win anticipation compared to neutral anticipation. Post hoc tests indicate that STs significantly increase BOLD activation from neutral to large win conditions but non-STs do not. STs marginally differed from non-STs during neutral but not during high win.

During the win-neutral anticipation contrasts, BOLD activations in the right amygdala showed a significant group by condition interaction (*F*_1,27_ = 9.04, *p* = .006, *η*^2^_p_ = 0.25, 90% CI = 0.05–0.46; Figure 3c), such that, in contrast to the IPL, STs exhibited a significant increase during the neutral condition relative to win activation (*t27* = −3.05, *p* = .025, Cohen’s d = −1.25, s.e. = 0.43, 95% CI = −2.12 – −0.37). These findings indicate salience-dependent modulation within the amygdala only for STs. In contrast to the IPL, non-STs did not differ between conditions or from STs during either condition in amygdala activation (*p*s > .1; see also Supplemental Figure S4 for additional sensitivity analysis after removing a possible outlier in the ST group). Taken together, the associations between cortical activation during salience processing for non-STs and subcortical activation during salience processing for STs, support two independent processes that contribute to the emergence of sign-tracking and non-sign-tracking behaviours. Whereas non-ST individuals showed greater activation in brain areas that have been associated with executive control, ST youth showed greater activation in the salience network.

To further understand the possible implications of these differential neural modulatory patterns, we used Pearson’s *r* to examine how percent signal change correspond to both externalizing behaviours and Pavlovian conditioned approach indices (Figure 4; Supplemental Table S6). Left IPL activation during win-neutral contrasts was negatively related to oppositional defiant problems (*r* = −0.43, *p* = .020), marginally negatively with negative affect (*r* = −0.34, *p* = .070), and marginally positively with inhibitory control (*r* = 0.36, *p* = .060), whereas left IPL during loss-neutral contrasts was negatively related to risk-taking behaviours (*r* = −0.45, *p* = .020). Although not all statistically significant, these associations support the notion that brain-behaviour associations might differ by phenotype. We further examined the correlations between percent signal change and PavCA scores. Activations in both regions during their respective contrasts were significantly correlated with the propensity to sign-track such that higher PavCA scores (sign-tracking) were related to less activity in the left IPL during both win-neutral (*r* = −0.46, *p* = .010) and loss-neutral (*r* = −0.45, *p* = .010) as well as greater activity in the amygdala during win-neutral (*r* = 0.48, *p* = .010). These findings are in agreement with the pre-clinical literature, suggesting that the behavioural endophenotype of sign-trackers is dominated by subcortical motivational systems; whereas that of non-sign-trackers is dominated by cortical processes.

### Environmental influences

Given that the stress response system directly influences both the dopaminergic reward system^51^ and animal sign- and goal tracking behaviours^8,52,53^, we also gathered information regarding environmental markers of adversity and protective factors including relationships and resources available to these youth.^54^ Importantly, STs reported lower household income (*t*_24.5_ = 2.81, *p* = .010, *d* = 0.94, s.e. = 0.39, 95% CI = 0.30–1.20), higher basic needs unaffordability (*t*_17.9_ = −2.28, *p* = .035, *d* = 0.79, s.e. = 0.42, 95% CI = −1.85 – −0.08), and lower markers of protective factors (*t*_33.7_ = 2.51, *p* = .017, *d* = 0.82, s.e. = 0.45, 95% CI = 0.22–2.06; Supplemental Table S3). Together, these findings support prior research addressing early life adversity as a potential influential driver for the divergence in phenotypic differences and associated vulnerabilities. While we did not have adequate power to do so, future studies with larger sample sizes would also benefit from examining the question whether social determinants of mental health are an important mediator or moderator for the expression of sign- or goal-tracking tendencies.

## Discussion

This study translated a paradigm previously developed to examine sign- and goal-tracking behaviours in rodents in order to determine whether these behavioural distinctions (1) could be observed in human youth, and are associated with (2) externalizing behaviours, or (3) reward-related neural activation patterns. First, we present evidence for sign-tracking behaviours in pre-adolescents. Second, STs were distinctive in both externalizing traits and neurobiological patterns consistent with impulse control disorders. Specifically, STs showed greater externalizing characteristics and salience-dependent subcortical reactivity to reward cues, whereas non-STs had fewer externalizing characteristics and more actively engaged cortical control regions. Third, both externalizing traits and the degree of sign-tracking behaviour correlated with neural modulation patterns such that those with higher BOLD activation in the left IPL also reportedly had fewer externalizing characteristics and had lower PavCA scores (non-STs) whereas those with increased BOLD activation in the amygdala also had higher PavCA scores (STs). Finally, those displaying a higher degree of sign-tracking behaviour also reported increased potential for early life stress (lower income, increased basic needs unaffordability, and fewer protective factors). Taken together, the phenotypic differences in our sample are consistent with rodent models of sign-tracking and support prior findings from humans with impulse control disorders.^5^

The differences in parent-reported behavioural tendencies between STs and non-STs in our sample reflect characteristic patterns in animal studies that are mechanistically tied to dual-systems processing.^5^ A fundamental characteristic of an incentive stimulus is its ability to bias attention and elicit approach behaviours suggestive of distinct cognitive/attentional control tendencies specific to this sign-trackers. In humans, attentional capture to reward-paired stimuli (resonant of sign-tracking behaviours) appears to be stronger for those with low cognitive control^55^ and directly linked to both impulsivity^56^ and substance use.^55^ Thus, the presence of increased attentional and executive control problems in STs, in conjunction with cortical activation patterns, suggests that, the attribution of incentive salience to reward-paired cues is driven by inefficient top-down cognitive and executive control.^19^ The other symptoms reported here for STs reflect patterns of behaviour consistent with the profile of dopamine-driven cue-reactivity in externalizing disorders, namely, defiance, aggression, and impulsivity. Notably, lower activation in cortical regions (i.e., in STs) is also correlated with behavioural reports of increased risk taking, defiance, and to a lesser extent externalizing behaviours, aggression, and lower inhibitory control. Finally, STs were reported to have increased fear-related behaviours, a finding that is supported by rodent models.^44,45^ This supports our finding that non-STs are not only reported to have fewer fear-related behaviours but also a neural modulatory profile indicative of emotion regulation and cognitive control.

Animal studies have shown a consistent double dissociation between top-down dominant cortical systems in GTs and subcortical cue-driven systems that dominate in STs.^14,57^ Here, we found concurring evidence in humans that, in contrast to STs, non-STs cognitively modulate their anticipatory responses in the IPL (and consistently across multiple cognitive control regions) but not the amygdala according to cue salience. The IPL has been implicated in probability and reward-related decision making^58^ and this pattern for non-STs may reflect cognitive discrimination in anticipatory and preparatory responses according to the salience of the cues. Whereas for STs, the non-modulation in the IPL appears to indicate indiscriminate cortical activation according to cue salience, suggesting that, cognitively, STs interpret all cues as salient and respond relatively equally across conditions. Thus, the selective modulation in this region by non-STs demonstrates distinct cognitive assessment of the cue and preparatory responses selectively applied to highly salient cues. STs do, however, affectively modulate subcortical (amygdala) activity in preparation for reward-paired cues. The amygdala contributes to contextual appraisal of rewards and motivational significance of incentives^15^ and is integral to neural processing of reward and reward cues.^59^ Therefore, the apparent reliance for STs on primarily subcortical structures to modulate responses and the fact that this salience-dependent modulation does not translate to cognitive action, supports a theory of increased vulnerability to impulse control disorders based on both individual differences in incentive salience attribution and mechanistic differences in reward processing. Our findings are also consistent with a recent fMRI analysis of model-based vs model-free learning in humans demonstrating that increased incentive salience attribution was associated with neural profiles of striatal reward prediction error signals in STs; whereas stronger cortical signals of state-prediction errors were evident in GTs.^31^ Together, STs in our sample are categorised both by behavioural Pavlovian responses to reward-paired cues and selectively modulated amygdala activity which points to a likelihood that these patterns are dopamine-dependent and supports previous pre-clinical animal reports of a dual-systems approach.

The dynamic nature of cortical development, top-down control, and neural organisation in pre-adolescence^37^ highlights the importance of considering developmental trajectories in light of both sign-tracking and externalizing tendencies. Specifically, adolescence is characterised by underdeveloped cortical downregulation of the amygdala that contributes to attentional/executive control deficits.^37,60^ It is likely, therefore, that this developmental variability is, at least in part, impacting both the neuromodulatory patterns and the significant skew in the tendency to sign-track. For instance, the limited goal-tracking behaviours is a notable deviance from rodent models. While age did not significantly differ between phenotypes, given our sample constraints, examination of these traits in larger samples with varying age ranges would help to further elucidate whether this is an effect of developmental stage or another factor not accounted for in our translation of this paradigm. However, in the absence of a significant age effect in PavCA behaviours in our sample, as well as statistical controls for age in place in neuroimaging models, it remains striking that we measured marked individual differences in both brain and behaviour. Further, the dynamic nature of both neural and behavioural traits during this age range, underscores the importance of measuring potential predictors of risk for psychopathology, as these characteristics may still be malleable. While more research is necessary to further clarify the developmental trajectory of sign- and goal-tracking and its impact on future psychopathology, measuring and characterising these phenotypes early and longitudinally as they evolve throughout development, underscores the potential utility of this paradigm to delineate possible preventative and interventive measures.

The individual differences in both the degree of sign-tracking and the development of impulse control deficits suggests the possibility of additional risk factors at play. Both sign-tracking^53^ and impulse control disorders^61^ are influenced by stressful early environments. Our finding that STs have lower income households, increased difficulty for affording basic needs, and less social support suggests probable environmental impacts on individual differences in sign-tracking tendencies and the underlying neurobiological mechanisms. The sensitivity of amygdala-frontal development in adolescence to stressful environmental input can bias the reward system to be more reactive to cues^62^ and vulnerable to reward-seeking and substance use.^61,63,64^ The finding that indicators of stressful environments are present more prominently in STs is consistent with this prior work and is supported by the corresponding psychopathological and neural profiles we observe in STs. However, this finding will need to be examined more fully in larger, more demographically diverse samples.

As a whole, these results provide evidence for sign-tracking in youth and a general neuromodulatory and behavioural profile consistent with externalizing traits; however, a number of limitations should be noted. First, our sample size was relatively small and demographically homogenous which limited our variance and power to perform multivariate analyses or complex neuroimaging models. In particular, future, higher powered studies would benefit from statistically addressing environmental factors including income and family history of psychopathology and substance use as well as individual-level factors such as pubertal stage to further account for possible sex differences. Additionally, a larger cohort would provide the opportunity to address the more limited extremes of sign- and goal-tracking behaviours and further validate if these neural and behavioural distinctions reflect differences in risk for externalizing disorders. Also of note is our use of monetary rewards for both the PavCA paradigm and the MID task which could influence salience for those participants coming from lower income households. Finally, additional methods of measurement should be considered (e.g., eye tracking, approach behaviours) that could further elucidate whether goal-tracking is, in fact, limited in pre-adolescence, or simply not adequately measured in this sample.

The current data strongly support the feasibility and utility of the sign-tracker/goal-tracker model of reward processing as a useful experimental approach and construct to measure individual differences in the propensity to attribute incentive salience to reward cues in youth. Moreover, these findings reveal a consistent and substantial pattern of neuromodulatory and externalizing trait responses that reliably dissociate bottom-up processing in STs from top-down cognitive control in non-STs. These results provide evidence in human youth for an underlying dual-systems mechanism of neural reward processing between cortical and subcortical control and link the degree of sign-tracking to externalizing aspects of psychopathology that may predispose some individuals to the development of impulse control disorders. The feasibility of this paradigm and the initial delineation of the circuitry associated with the degree of sign-tracking is enhanced by the extensive knowledge base obtained from prior animal studies, which provide neuroscience-based rationale for future studies elucidating the underlying mechanisms of risk for externalizing psychopathology. While there is still more to be discovered, particularly regarding the developmental trajectory, stability, or malleability of these phenotypes in humans, these data offer promise in identifying sign- and goal-tracking phenotypes in humans and present initial evidence detailing corresponding neural profiles and behavioural traits consistent with indicators of impulse control disorders, helping to pave the way for future research with clinical applications.

## Methods

### Participants and Sample Selection

The sample consisted of 9-12-year-old children (N = 40, *m* = 9.6 years, *sd* = 0.93; Table 1). All tasks and measures were used in the whole sample. Participants were excluded if they received a diagnosis of a severe learning disorder, Axis 1 psychiatric disorder, Attention Deficit/Hyperactivity Disorder or Pervasive Developmental Disorder (e.g., Autism Spectrum Disorder) as these conditions may affect attentional control and possibly bias the measurement of attention to each stimulus, or if they endorsed MRI contraindications including non-correctable vision, hearing, or sensorimotor impairments, claustrophobia, large body size, or irremovable ferromagnetic metal implements or dental appliances. Participants were recruited via fliers and online advertisements. Youth participated with one parent/caregiver present and all participants were compensated for their time. Caregivers completed questionnaires including demographics, youth behaviours, and family environment. Youth completed the sign- and goal-tracking task at the beginning of each session followed by self-report questionnaires, behavioural and neurocognitive tasks, practice tasks for imaging sessions, and neuroimaging. All study procedures were approved by an institutional review board and participants provided informed consent. A power analysis was used to estimate the appropriate sample size based on the index of Pavlovian conditioned behaviour found in an existing study of sign-tracking and goal-tracking in humans.^30^ An *a priori* power analysis using G*Power software^65^ indicated that a sample of 40 would be sufficient to detect a large effect with 80% power (alpha=.05, two-tailed) for independent samples *t*-tests using two groups (ST and non-ST). While likely under-powered for more complex analyses involving additional covariates, these results will be critical for estimating effect sizes for future research.

In addition to validation of the sign-tracker/goal-tracker paradigm, the purpose of this study was to identify behaviours and neurological profiles associated with sign-tracking phenotypes. The age range for this sample was selected for multiple reasons. First, the average age of onset for externalizing disorders is 11 years old and many externalizing symptoms emerge during this age range.^2^ Second, symptoms of a range of psychiatric conditions (e.g., depression, anxiety, substance use) typically begin to emerge later in adolescence^66^, making pre-adolescence a prime target for mental health screening and development of preventative interventions. Further, the timing of prefrontal cortex development, responsible for decision-making and higher-order executive functions^37^, pre-adolescent youth may be more likely to engage in behaviours influenced by enhanced motivational drive^67,68^ than adults, likely impacting the detection of sign-tracking or goal-tracking phenotypes in humans. And finally, we used this age range in an attempt to maximise the quality of neuroimaging data collection in children.

## Materials

### Sign and Goal Tracking: Apparatus and Paradigm Development

The Pavlovian conditioning apparatus used is described in Joyner et al. (2018) and was built to mimic the animal model as closely as possible. The apparatus consisted of two solid-coloured response boxes, built to look like building blocks and be appealing to youth but not inherently rewarding. The boxes were: (1) the CS box containing a lever, which illuminated and extended from the box, and (2) the US box containing a small metal tray into which the reward was dispensed. In the CS box, a linear actuator, consisting of a 12V DC motor (Bühler Motor, GmbH, Nuremberg, Germany, www.buehlermotor.com) and a worm drive, was used to extend and retract the lever. In the US box, a reward dispenser with infrared sentry (Med Associates, Inc, Fairfax, VT, www.medassociates.com) was used to release the reward. A touch sensor, based on a field-effect transistor, was used to detect a participant’s touches to the metal reward tray. The hardware system was operated via an Arduino UNO microcontroller (Arduino, LLC, Somerville, MA, www.arduino.cc). The microcontroller was programmed to provide three output signals, controlling the lever movement, the lever light, and the reward dispenser. Two Arduino inputs were programmed to record the lever presses and the reward tray touches. The experimental protocol was implemented in custom software written in MATLAB (MathWorks, Inc, Natick, MA, www.mathworks.com). The code was run by a researcher on a laptop, communicating with the Arduino via a USB connection. Response times (in ms) corresponding to all lever presses and reward tray touches in each trial were saved to a file. The CS and US boxes were spaced approximately 12 inches apart to reduce the likelihood of participants simultaneously engaging with both. The left/right positioning of the boxes was counterbalanced between participants to minimise lateral bias.

The structure of the Pavlovian conditioning paradigm was primarily modeled after rodent studies described by Flagel and colleagues (2008).^69^ This paradigm consists of multiple response-independent trials during which a lever (CS) extends, and upon retraction, a reward portion (US) is dispensed. A randomly selected inter-trial interval (ITI) follows each trial, after which the next trial begins. In typical animal models^69^, conditioning sessions consist of 25 trials lasting 30-45 minutes and occur across multiple days. Modeled after Joyner and colleagues (2018)^36^, we condensed training into a single session to make this more feasible for our population and avoid participant burden (e.g., multiple trips to the lab). Within this session, we conducted four blocks of ten trials each, with the total session lasting approximately 20-30 minutes. Each trial consisted of the lever illuminating and extending for 8.3s, and upon retraction, the token reward (US) was dispensed into the tray. The ITI period was programmed to last for a randomly selected time, either 8, 16, 24, or 32s. Each block was followed by a “wiggle break” lasting up to 45 seconds, during which the child was given the opportunity to relax and reset before the next trial. This setup was intended to maximise participant attention, minimise fatigue, while retaining enough length in the session for associative learning to occur. In order to capture number of and latency to contacts to each stimulus, we measured these behaviours in line with the traditional measurement of rodent sign- and goal-tracking, the MATLAB program controlling the apparatus recorded the number and timing of contacts to the CS lever and US reward tray during CS presentation and ITI (Figure 1).

To assess learning of conditioned responses to the CS, we examined behavioural responses to the CS using two-sided linear mixed effects models with the factors time (Block 1-4), phase (CS, ITI), and phenotype (ST, non-ST) for lever- and reward-directed behaviours (Supplemental Figure S2) and PavCA (Figure 1c). These models included age and sex as covariates and a random intercept for subject. For post hoc tests we used estimated marginal means. We investigated behaviours both during the CS periods (as seen in animal models) and ITI periods because, unlike animal models, our sample displayed lever-directed behaviours, albeit low levels, during the ITI-period (i.e., when the lever was retracted; CS lever contacts, *m* = 4.06, *sd* = 5.73; ITI lever contacts, *m* = 0.59, *sd* = 1.53). This may indicate an effect of investigative or exploratory behaviours common during this developmental stage^70^ or limited attentional capacity during the longer ITIs and will be an important methodological consideration in human translation of this paradigm in future studies. Further, because human subjects attempted to interact with the lever even during the ITI (i.e. when it was retracted), and because CS lever contacts correlated positively with ITI lever contacts (*r* = 0.59, *p* < .001, Supplemental Table S2), a normalised frequency score was calculated for lever- and reward-directed behaviours (contacts and probability) in order to adequately compare between CS and ITI periods. To normalise, we divided each measure by the length of their respective phase. A non-normalised score was used for PavCA scores in the main text.

When determining an appropriate reward to use as the US, we took several factors into consideration. First, pilot testing primary food rewards in this age group (chocolate candies, as in Joyner et al.^36^) demonstrated very minimal incentive toward the reward. Therefore, in order to choose an item that would be (1) rewarding to the study population and (2) small and consistent in shape to both adequately dispense from the machine and reflect the physical characteristics of the candies used in Joyner et al.,^36^ we ultimately decided to use colourful wooden beads. Participants were informed these tokens were worth $0.20 cents each and could be exchanged for their choice of prizes or money (totaling $8) at the end of the session. The use of a secondary reinforcement rather than primary is a notable deviation from rodent models and Joyner and colleagues^36^, however, we believe the benefit of increased participant engagement and reward motivation made this justifiable.

### Measures

#### Parent-reported psychopathology

Two parent-report measures that have been developmentally validated for 9-12-year-olds were used to identify youth psychopathology symptoms. First, the Early Adolescent Temperament Questionnaire (EATQ-R)^41^ is a 62-item parent-report measure of youth temperament and self-regulation in 9-15-year-olds. Subscales include effortful control (measures of attention, inhibitory control, and activation control), surgency (measures of surgency, fear, and shyness), negative affect (measures of aggression, frustration, and depressive mood), and affiliativeness. Second, the Child Behaviour Checklist (CBCL)^40^ measures dimensional psychopathology and is normed by age, sex, and ethnicity. It is a parent-reported youth behavioural questionnaire that includes eight empirically-based subscales for anxious/depressed, withdrawn/depressed, somatic complaints, social problems, thought problems, attention problems, rule breaking, and aggression as well as DSM-oriented subscales for attention deficit-hyperactivity, affective, anxiety, somatic, conduct, and oppositional defiant problems.

#### Family Demographics and Functioning

Parents responded to demographic questions from the PhenX survey toolkit^71,72^ including household income and 7 items addressing economic adversity and basic needs unaffordability. Additional environmental influences included in our analyses were markers of family environment including the Protective and Compensatory Experiences Scale (PACEs)^54^, a 10-item parent-report scale measuring factors contributing to resiliency in childhood including relationships and resources available to youth. Finally, parents reported on the family environment with the 90-item Family Environment Scale.^73,74^ Items were true/false and subscales for cohesion, organisation, recreational activities, and conflict were included.

#### Youth self-report and neurocognitive functioning

Youth reported impulsive and inhibitory behaviours using the Urgency, Premeditation, Perseverance, Sensation Seeking, Positive Urgency (UPPS-P), Impulsive Behaviour Scale and the behavioural inhibition system/ behavioural approach system (BIS/BAS)^75^ scale. A modified version of the UPPS-P from PhenX for children was used^76,77^ consisting of 20 self-report questions addressing youth impulsive behaviours including subscales for negative and positive urgency, lack of premeditation and perseverance, and sensation seeking. An abridged version of the BIS/BAS scale was used including 24 items with subscales for drive, fun seeking, reward responsiveness, and inhibition.^75,77^ Youth neurocognitive functioning was measured using the National Institutes for Health Toolbox Neurocognitive Battery specified for ages 7-17.^78,79^ All tests were administered measuring executive functioning, episodic memory, language, processing speed, working memory, and attention. For analysis purposes we included the age-corrected composite scores of fluid and crystalized cognitive functioning.

### Neuroimaging

#### Head Motion Prevention

To prevent and minimise motion: (1) participants watched an age-appropriate informational video explaining MRI safety and the importance of staying still; (2) prior to the scan session, participants completed motion compliance training in mock scanners using head motion detection/feedback; (3) the head was stabilised in head coils; and (4) the MR technologist modeled relaxation techniques. Youth vision was screened, and MRI-safe corrective lenses were provided if necessary.

#### Monetary Incentive Delay (MID) fMRI Task

Participants completed two runs of the Monetary Incentive Delay (MID) task.^50^ Each trial began with a cue presented for 2 seconds indicating the trial type, with a pink circle indicating a potential gain of $5 or $0.20, a blue triangle indicating no gain or loss, and a yellow square indicating a potential loss of $5 or $0.20 so that there were 5 trial types: high-loss, low-loss, neutral, low-win, and high-win. A 1.5 to 4 second fixation followed the cue, and then a black target was presented. Participants were instructed to press a button while the target was on the screen, with target duration varying between 0.15 and 0.5s. On gain trials, participants were rewarded for successfully hitting the target and would neither gain nor lose money for missing it. On loss trials, participants lost the indicated amount when they missed and neither gained nor lost when they hit the target. The outcome of each trial was presented immediately after each response and lasted 2 seconds minus the duration of the target. Target duration was initialised based on performance during a practice session completed outside the scanner and updated during scanning so that participants would succeed on approximately 60% of trials, leading to mean earnings of $20 with a maximum of $60. Each run contained 50 trials (10 of each trial type) and lasted 5 minutes 42 seconds (Supplemental Figure S3).

BOLD imaging during the MID task took place on two identical GE MR750 3T scanners using multiband acquisition with an acceleration factor of 6 and the following parameters: 60 axial slices, TR/TE = 800/30ms, FOV/slice = 216/2.4mm, 90 x 90 matrix producing 2.4mm isotropic voxels, 419 volumes for 5 minutes 35s of scan time per run. Additionally, high-resolution structural images were obtained through a 3D sagittal T1-weighted magnetization-prepared rapid acquisition with gradient echo sequence (TR/TE = 6/2.92 ms, FOV/slice = 256 × 256/1mm, 208 sagittal slices).

#### fMRI Data Preprocessing

fMRI data were preprocessed using the Analysis of Functional Neuroimaging (AFNI, http://anfi.nimh.nih.gov) software.^80^ Steps included removal of the first 10 volumes to allow for signal stabilization, despiking, slice timing correction, co-registration with the anatomical volume, motion correction, non-linear warp to MNI space with resampling to 2mm isotropic voxels, scaling to percent signal change, and application of a 4mm Gaussian FWHM smoothing kernel. A general linear model was applied with regressors for the six motion parameters, three polynomial terms, and 2-second block regressors for each of the five trial types. Censoring was applied at the regression step so that any TRs with the Euclidean norm of the six motion parameter derivatives greater than 0.3 were removed, along with TRs where greater than 10 percent of brain voxels were outliers (estimated with 3dToutcount). The estimated beta coefficients from this single subject analysis were taken to the group level and are interpreted in terms of percent signal change for each condition.

### Statistical Analysis

Analyses were conducted in the R System for Statistical Computing.^81^ Feasibility of the sign- and goal-tracking task was assessed in multiple ways. First, frequency counts were used to identify qualitative and quantitative levels of child engagement in the paradigm and how well the participants tolerated the task. Second, we assessed behavioural responses during CS and ITI periods using means and standard deviations of each behaviour by block. Additionally, Pearson *r* correlations using the psych package in R^82^ were used within behaviours to verify measurement of lever- or reward-directed behaviours. Blocks 1 and 2 were identified as a training phase, therefore the behaviours from Blocks 3 and 4 were averaged for each PavCA index score. To classify categorical phenotypic groups, we used a PavCA value of 0.5 or greater to define sign-tracking and less then −0.5 to define goal-tracking, consistent with categorisation used in animal models.^83^ Behavioural differences by phenotype were assessed using two-sided Welch two sample *t*-tests in the R package stats.^81^ Finally, we assessed the demonstration of learning a conditioned response to the CS by examining behavioural responses to the CS over all four Blocks (40 trials total). We used linear mixed effects models using the R package lme4^84^ to test for three-way interactions between time (Block 1-4), phase (CS, ITI), and phenotype (ST, non-ST) for lever- and reward-directed behaviours. The R package emmeans^85^ was used to assess planned post hoc comparisons. Error bars in figures represent standard error of the mean calculated in the Rmisc^86^ package in R.

Recent research using animal models has shown that ITI duration may also impact the likelihood of displaying each CR and sign-tracking behaviour appears to be more likely during a longer ITI periods^87^ due to a weakened association between contacting the location of the US and receiving a reward. Additionally, the responses during ITI periods in our sample were relatively high compared to typical animal behaviours, therefore, to adequately compare between CS and ITI periods, scores were normalised by dividing each score by the length of each respective phase (8 seconds for CS, and 8, 16, 24, or 32 seconds for each ITI).

#### Behavioural outcome measures

We conducted Welsh independent samples *t*-tests using the R package stats to examine phenotypic differences in symptoms and environmental variables. Tests were *p*-value corrected for multiple comparisons using the FDR corrections. All variables were tested for normality and transformed using the optLog package in R where necessary. CBCL subscales were all log transformed.

#### Whole Brain Analyses

We performed a whole-brain voxelwise linear mixed effects analysis using AFNI’s 3dLME^88^ with fixed effects for group, condition, age, sex, a group by condition interaction and a random intercept for subject. We followed this with planned contrasts investigating a group (ST, non-ST) by condition (high-win vs neutral or high-loss vs neutral) interaction. 3dFWHMx (with the newer -acf option) was used to estimate the smoothness of the residuals of the group model, and this smoothness was used along with 3dClustSim to perform cluster-wise correction. Significant clusters are reported using a voxelwise *p*-value threshold of 0.005 and α < 0.05 at the cluster level (N= 11.73 voxels).

---

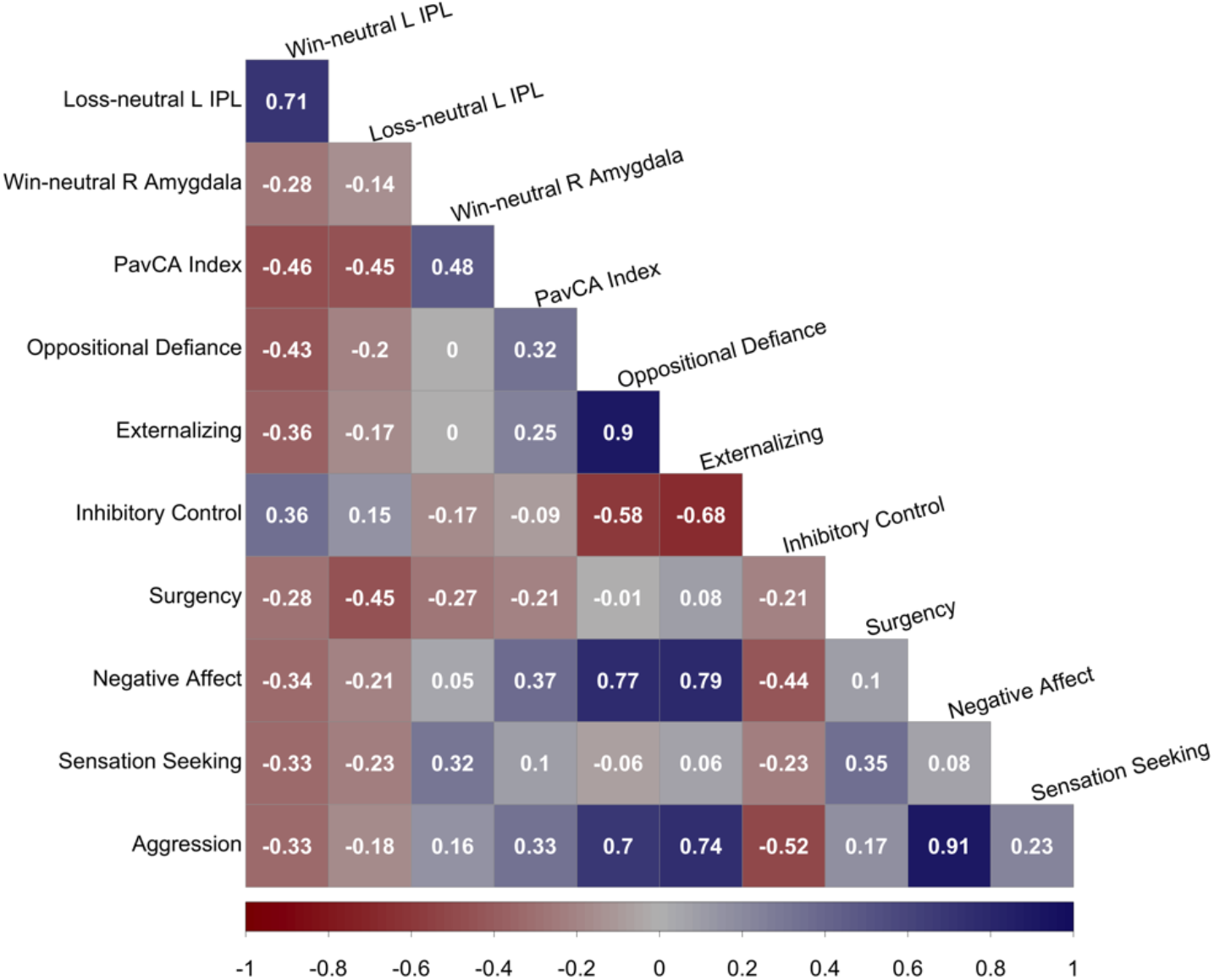

## Acknowledgements

This work is supported by the Centers of Biomedical Research Excellence (CoBRE) grant (P20 GM121312, PI: M. Paulus) funded by the National Institutes of Health and the William K. Warren Foundation. S.B. Flagel’s effort on this project was funded, in part, by the Pritzker Neuropsychiatric Disorders Research Consortium Fund LLC (http://www.pritzkerneuropsych.org). We thank J. J. Colaizzi for graphical design assistance, J. Touthang for help with data processing and technical assistance, and E. Pribil for data collection. The funders had no role in the study design, data collection or analysis, decision to publish, or preparation of the manuscript.

## Author Contributions

J.M.C., S.B.F., A.N.G., M.B., and M.P.P. designed the study. J.M.C., G.C., J.C., T.A. conducted pilot tests, feasibility tests, and study setup. G.C., J.C., T.A. acquired the data with supervision from J.M.C. and M.P.P. V.Z. provided technical and mechanical support. J.M.C. analysed the data with supervision and input from S.B.F., A.N.G., R.K., and M.P.P. J.M.C. wrote the manuscript. All authors read and revised the manuscript and provided critical intellectual contributions.

## Additional information

Supplementary materials are available for this paper

## Referecnes

1. Castellanos-Ryan N, Struve M, Whelan R, Banaschewski T, Barker GJ, Bokde ALW. Neural and cognitive correlates of the common and specific variance across externalizing problems in young adolescence. The American Journal of Psychiatry. 2014;171(12):1310–1319.

2. Merikangas KR, He JP, Burstein M. Lifetime prevalence of mental disorders in U.S. adolescents: results from the National Comorbidity Survey Replication--Adolescent Supplement (NCS-A). Journal of the American Academy of Child and Adolescent Psychiatry. 2010;49(10):980–989.

3. Biederman J, Petty CR, Dolan Cea. The long-term longitudinal course of oppositional defiant disorder and conduct disorder in ADHD boys: findings from a controlled 10-year prospective longitudinal follow-up study. Psychological Medicine. 2008;38(7):1027–1036.

4. Saunders BT, Robinson TE. Individual variation in resisting temptation: implications for addiction. Neuroscience & Biobehavioral Reviews. 2013;37(9 Pt A):1955–1975.

5. Colaizzi JM, Flagel SB, Joyner MA, Gearhardt AN, Stewart JL, Paulus MP. Mapping sign-tracking and goal-tracking onto human behaviors. Neuroscience & Biobehavioral Reviews. 2020;111:84–94.

6. Phillips KB, Sarter M. Addiction vulnerability and the processing of significant cues: sign-, but not goal-, tracker perceptual sensitivity relies on cue salience. Behavioral Neuroscience. 2020;134(2):133–143.

7. Robinson TE, Carr C, Kawa AB. The propensity to attribute incentive salience to drug cues and poor cognitive control combine to render sign-trackers susceptible to addiction. In: A T, JD M, eds. Sign-tracking and drug addiction. Ann Arbor: Maize Books, Michigan Publishing; 2018.

8. Flagel SB, Akil H, Robinson TE. Individual differences in the attribution of incentive salience to reward-related cues: Implications for addiction. Neuropharmacology. 2009;56 Suppl 1:139–148.

9. Robinson TE, Flagel SB. Dissociating the predictive and incentive motivational properties of reward-related cues through the study of individual differences. Biological Psychiatry. 2009;65(10):869–873.

10. Robinson TE, Yager LM, Cogan ES, Saunders BT. On the motivational properties of reward cues: Individual differences. Neuropharmacology. 2014;76 Pt B:450–459.

11. Robinson TE, Flagel SB. Dissociating the predictive and incentive motivational properties of reward-related cues through the study of individual differences. Biological psychiatry. 2009;65(10):869–873.

12. Boakes RA. Performance on learning to associate a stimulus with positive reinforcement. In: Davis H, Hurwitz H, eds. Operant-Pavlovian Interactions. Hillsdale, NJ: Erlbaum; 1977:67–97.

13. Hearst E, Jenkins H. Sign-tracking: the Stimulus-reinforcer Relation and Directed Action. Austin 1974.

14. Sarter M, Phillips KB. The neuroscience of cognitive-motivational styles: sign-and goal-trackers as animal models. Behavioral Neuroscience. 2018;132(1):1–12.

15. Baxter MG, Murray EA. The amygdala and reward. Nature Reviews Neuroscience. 2002;3(7):563–573.

16. Kalivas PW, Volkow ND. The neural basis of addiction: a pathology of motivation and choice. The American Journal of Psychiatry. 2005;162(8):1403–1413.

17. Flagel SB, Clark JJ, Robinson TE, et al. A selective role for dopamine in stimulus-reward learning. Nature. 2011;469(7328):53–57.

18. Flagel SB, Robinson TE. Neurobiological basis of individual variation in stimulus-reward learning. Current Opinion in Behavioral Sciences. 2017;13:178–185.

19. Koshy Cherian A, Kucinski A, Pitchers K, et al. Unresponsive choline transporter as a trait neuromarker and a causal mediator of bottom-up attentional biases. The Journal of Neuroscience. 2017;37(11):2947–2959.

20. Lovic V, Keen D, Fletcher PJ, Fleming AS. Early-life maternal separation and social isolation produce an increase in impulsive action but not impulsive choice. Behavioral Neuroscience. 2011;125(4):481–491.

21. Paolone G, Angelakos CC, Meyer PJ, Robinson TE, Sarter M. Cholinergic control over attention in rats prone to attribute incentive salience to reward cues. The Journal of Neuroscience. 2013;33(19):8321–8335.

22. Pitchers KK, Phillips KB, Jones JL, Robinson TE, Sarter M. Diverse roads to relapse: a discriminative cue signaling cocaine availability is more effective in renewing cocaine seeking in goal trackers than sign trackers and depends on basal forebrain cholinergic activity. The Journal of Neuroscience. 2017;37(30):7198–7208.

23. Saunders BT, Robinson TE. The role of dopamine in the accumbens core in the expression of Pavlovian-conditioned responses. European Journal of Neuroscience. 2012;36(4):2521–2532.

24. Sarter M, Paolone G. Deficits in attentional control: cholinergic mechanisms and circuitry-based treatment approaches. Behavioral Neuroscience. 2011;125(6):825–835.

25. Moeller SJ, Paulus MP. Toward biomarkers of the addicted human brain: Using neuroimaging to predict relapse and sustained abstinence in substance use disorder. Progress in Neuro-Psychopharmacology & Biological Psychiatry. 2018;80(Pt B):143–154.

26. Volkow ND, Wang GJ, Fowler JS, Tomasi D, Telang F, Baler R. Addiction: decreased reward sensitivity and increased expectation sensitivity conspire to overwhelm the brain’s control circuit. Bioessays. 2010;32(9):748–755.

27. Cope LM, Martz ME, Hardee JE, Zucker RA, Heitzeg MM. Reward activation in childhood predicts adolescent substance use initiation in a high-risk sample. Drug and Alcohol Dependence. 2019;194:318–325.

28. Anderson BA, Faulkner ML, Rilee JJ, Yantis S, Marvel CL. Attentional bias for nondrug reward is magnified in addiction. Experimental and Clinical Psychopharmacology. 2013;21(6):499–506.

29. Stautz K, Cooper A. Impulsivity-related personality traits and adolescent alcohol use: a meta-analytic review. Clinical Psychology Review. 2013;33(4):574–592.

30. Garofalo S, di Pellegrino G. Individual differences in the influence of task-irrelevant Pavlovian cues on human behavior. Frontiers in Behavioral Neuroscience. 2015;9:163.

31. Schad DJ, Rapp MA, Garbusow M, et al. Dissociating neural learning signals in human sign-and goal-trackers. Nature Human Behaviour. 2019;4(2):201–214.

32. Failing M, Nissens T, Pearson D, Le Pelley M, Theeuwes J. Oculomotor capture by stimuli that signal the availability of reward. Journal of Neurophysiology. 2015;114(4):2316–2327.

33. Le Pelley ME, Pearson D, Griffiths O, Beesley T. When goals conflict with values: counterproductive attentional and oculomotor capture by reward-related stimuli. Journal of Experimental Psychology: General. 2015;144(1):158–171.

34. Pearson D, Donkin C, Tran SC, Most SB, Le Pelley ME. Cognitive control and counterproductive oculomotor capture by reward-related stimuli. Visual Cognition. 2015;23(1-2):41–66.

35. Pearson D, Osborn R, Whitford TJ, Failing M, Theeuwes J, Le Pelley ME. Value-modulated oculomotor capture by task-irrelevant stimuli is a consequence of early competition on the saccade map. Attention, Perception, & Psychophysics. 2016;78(7):2226–2240.

36. Joyner MA, Gearhardt AN, Flagel SB. A translational model to assess sign-tracking and goal-tracking behavior in children. Neuropsychopharmacology. 2018;43(1):228–229.

37. Casey BJ, Jones RM. Neurobiology of the adolescent brain and behavior. Journal of the American Academy of Child and Adolescent Psychiatry. 2010;49(12):1189–1201.

38. Flagel SB, Watson SJ, Akil H, Robinson TE. Individual differences in the attribution of incentive salience to a reward-related cue: influence on cocaine sensitization. Behav Brain Res. 2008;186(1):48–56.

39. Meyer PJ, Lovic V, Saunders BT, et al. Quantifying individual variation in the propensity to attribute incentive salience to reward cues. PLOS One. 2012;7(6):e38987.

40. Achenbach TM. The Achenbach System of Empirically Based Assessemnt (ASEBA): Development, Findings, Theory, and Applications. Burlington, VT: University of Vermont Research Center for Children, Youth, & Families; 2009.

41. Capaldi DM, Rothbart MK. Development and validation of an early adolescent temperament measure. Journal of Early Adolescence. 1992;12:153–173.

42. Frodl T. Comorbidity of ADHD and Substance Use Disorder (SUD): a neuroimaging perspective. Journal of Attention Disorders. 2010;14(2):109–120.

43. Matthews M, Nigg JT, Fair DA. Attention deficit hyperactivity disorder. Current Topics in Behavioral Neurosciences. 2014;16:235–266.

44. Morrow JD, Saunders BT, Maren S, Robinson TE. Sign-tracking to an appetitive cue predicts incubation of conditioned fear in rats. Behavioural Brain Research. 2015;276:59–66.

45. Maria-Rios CE, Morrow JD. Mechanisms of shared vulnerability to post-traumatic stress disorder and substance use disorders. Frontiers in Behavioral Neuroscience. 2020;14:6.

46. Flagel SB, Robinson TE, Clark JJ, et al. An animal model of genetic vulnerability to behavioral disinhibition and responsiveness to reward-related cues: implications for addiction. Neuropsychopharmacology. 2010;35(2):388–400.

47. Verdejo-Garcia A, Bechara A, Recknor EC, Perez-Garcia M. Negative emotion-driven impulsivity predicts substance dependence problems. Drug and Alcohol Dependence. 2007;91(2-3):213–219.

48. Noordermeer SD, Luman M, Oosterlaan J. A systematic review and meta-analysis of neuroimaging in Oppositional Defiant Disorder (ODD) and Conduct Disorder (CD) taking Attention-Deficit Hyperactivity Disorder (ADHD) into account. Neuropsychology Review. 2016;26(1):44–72.

49. Tomie A, Morrow JD. The role of sign-tracking in drug addiction. Ann Arbor, MI.: Michigan Publishing; 2018.

50. Knutson B, Westdorp A, Kaiser E, Hommer D. FMRI visualization of brain activity during a monetary incentive delay task. Neuroimage. 2000;12(1):20–27.

51. Piazza PV, Le Moal ML. Pathophysiological basis of vulnerability to drug abuse: role of an interaction between stress, glucocorticoids, and dopaminergic neurons. Annual Review of Pharmacology and Toxicology. 1996;36:359–378.

52. Beckmann JS, Bardo MT. Environmental enrichment reduces attribution of incentive salience to a food-associated stimulus. Behavioural Brain Research. 2012;226(1):331–334.

53. Lomanowska AM, Lovic V, Rankine MJ, Mooney SJ, Robinson TE, Kraemer GW. Inadequate early social experience increases the incentive salience of reward-related cues in adulthood. Behavioural Brain Research. 2011;220(1):91–99.

54. Hays-Grudo J, Morris AS. Adverse and protective childhood experiences: a developmental perspective. Washington DC: APA Press; 2020.

55. Albertella L, Copeland J, Pearson D, Watson P, Wiers RW, Le Pelley ME. Selective attention moderates the relationship between attentional capture by signals of nondrug reward and illicit drug use. Drug and Alcohol Dependence. 2017;175:99–105.

56. Albertella L, Le Pelley ME, Chamberlain SR, Westbrook F, Fontenelle LF, R. S. Reward-related attentional capture is associated with severity of addictive and obsessivecompulsive behaviors. Psychology of Addictive Behaviors. 2019;33(5):495–502.

57. Pitchers KK, Kane LF, Kim Y, Robinson TE, Sarter M. ‘Hot’ vs. ‘cold’ behavioural-cognitive styles: motivational-dopaminergic vs. cognitive-cholinergic processing of a Pavlovian cocaine cue in sign-and goal-tracking rats. European Journal of Neuroscience. 2017;46(11):2768–2781.

58. Si R, Rowe JB, Zhang J. Functional localization and categorization of intentional decisions in humans: A meta-analysis of brain imaging studies. Neuroimage. 2021;242:118468.

59. Volkow ND, Fowler JS, Wang GJ. The addicted human brain viewed in the light of imaging studies: brain circuits and treatment strategies. Neuropharmacology. 2004;47:3–13.

60. Williams BR, Ponesse JS, Schachar RJ, Logan GD, Tannock R. Development of inhibitory control across the life span. Developmental Psychology. 1999;35(1):205–213.

61. Wakeford AGP, Morin EL, Bramlett SN, Howell LL, Sanchez MM. A review of nonhuman primate models of early life stress and adolescent drug abuse. Neurobiology of Stress. 2018;9:188–198.

62. Somerville LH, Jones RM, Casey BJ. A time of change: behavioral and neural correlates of adolescent sensitivity to appetitive and aversive environmental cues. Brain and Cognition. 2010;72(1):124–133.

63. Oswald LM, Wand GS, Kuwabara H, Wong DF, Zhu S, Brasic JR. History of childhood adversity is positively associated with ventral striatal dopamine responses to amphetamine. Psychopharmacology (Berl). 2014;231(12):2417–2433.

64. Roos LE, Horn S, Berkman ET, Pears K, Fisher PA. Leveraging translational neuroscience to inform early intervention and addiction prevention for children exposed to early life stress. Neurobiology of Stress. 2018;9:231–240.

65. Faul F, Erdfelder E, Lang AG, Buchner A. G*Power 3: a flexible statistical power analysis program for the social, behavioral, and biomedical sciences. Behavior Research Methods. 2007;39(2):175–191.

66. Bongers IL, Koot HM, J. vdE, Verhulst FC. Developmental trajectories of externalizing behaviors in childhood and adolescence. Child Development. 2004;74(5):1523–1537.

67. Rollins BY, Loken E, Savage JS, Birch LL. Measurement of food reinforcement in preschool children. Associations with food intake, BMI, and reward sensitivity. Appetite. 2014;72:21–27.

68. Silvers JA, Insel C, Powers A, et al. Curbing craving: behavioral and brain evidence that children regulate craving when instructed to do so but have higher baseline craving than adults. Psychol Sci. 2014;25(10):1932–1942.

69. Flagel SB, Watson SJ, Akil H, Robinson TE. Individual differences in the attribution of incentive salience to a reward-related cue: influence on cocaine sensitization. Behavioural Brain Research. 2008;186(1):48–56.

70. Berlyne DE. Curiosity and exploration. Science. 1966;153(3731):25–33.

71. Barch DM, Albaugh MD, Avenevoli S, et al. Demographic, physical and mental health assessments in the adolescent brain and cognitive development study: Rationale and description. Dev Cogn Neurosci. 2018;32:55–66.

72. Stover PJ, Harlan WR, Hammond JA, Hendershot T, Hamilton CM. PhenX: a toolkit for interdisciplinary genetics research. Curr Opin Lipidol. 2010;21(2):136–140.

73. Moos RH, Moos BS. Family Environment Scale Manual. Palo Alto, CA: Consulting Psychologists Press; 1994.

74. Zucker RA, Gonzalez R, Feldstein Ewing SW, et al. Assessment of culture and environment in the Adolescent Brain and Cognitive Development Study: Rationale, description of measures, and early data. Dev Cogn Neurosci. 2018;32:107–120.

75. Pagliaccio D, Luking KR, Anokhin AP, et al. Revising the BIS/BAS Scale to study development: measurement invariance and normative effects of age and sex from childhood through adulthood. Psychological Assessment. 2016;28(4):429–442.

76. Zapolski TC, Stairs AM, Settles RF, Combs JL, Smith GT. The measurement of dispositions to rash action in children. Assessment. 2010;17(1):116–125.

77. Barch DM, Albaugh MD, Avenevoli S, et al. Demographic, physical and mental health assessments in the adolescent brain and cognitive development study: Rationale and description. Developmental Cognitive Neuroscience. 2018;32:55–66.

78. Gershon RC, Wagster MV, Hendrie HC, Fox NA, Cook KF, Nowinski CJ. NIH toolbox for assessment of neurological and behavioral function. Neurology. 2013;12(80):(11 Suppl 13):S12–16.

79. Luciana M, Bjork JM, Nagel BJ, et al. Adolescent neurocognitive development and impacts of substance use: Overview of the adolescent brain cognitive development (ABCD) baseline neurocognition battery. Dev Cogn Neurosci. 2018;32:67–79.

80. Cox RW. AFNI: software for analysis and visualization of functional magnetic resonance neuroimages. Computers and Biomedical research. 1996;29(3):162–173.

81. R Core Team. R: a language and environment for statistical computing. Vienna, Austria 2020.

82. Revelle W. psych: procedures for psychological, psychometric, and personality research. In: Northwestern University E, Illinois, ed: R package version 2.1.9; 2021.

83. Yager LM, Pitchers KK, Flagel SB, Robinson TE. Individual variation in the motivational and neurobiological effects of an opioid cue. Neuropsychopharmacology : official publication of the American College of Neuropsychopharmacology. 2015;40(5):1269–1277.

84. Bates D, Maechler M, Bolker B, Walker S. Fitting linear mixed-effects models using lme4. Journal of Statistical Software. 2015;67(1):1–48.

85. emmeans: estimated marginal means, aka least-squares means. 2021. https://CRAN.R-project.org/package=emmeans.

86. Rmisc: Ryan Miscellaneous. 2013. https://CRAN.R-project.org/package=Rmisc.

87. Lee B, Gentry RN, Bissonette GB, et al. Manipulating the revision of reward value during the intertrial interval increases sign tracking and dopamine release. PLoS Biol. 2018;16(9):e2004015.

88. Chen G, Saad ZS, Britton JC, Pine DS, Cox RW. Linear mixed-effects modeling approach to fMRI group analysis. NeuroImage. 2013;73:176–190.

