## Supplemental Tables and Figures for "The propensity to sign-track is associated with externalizing behaviour and distinct patterns of reward-related brain activation in youth"

Supplemental Material

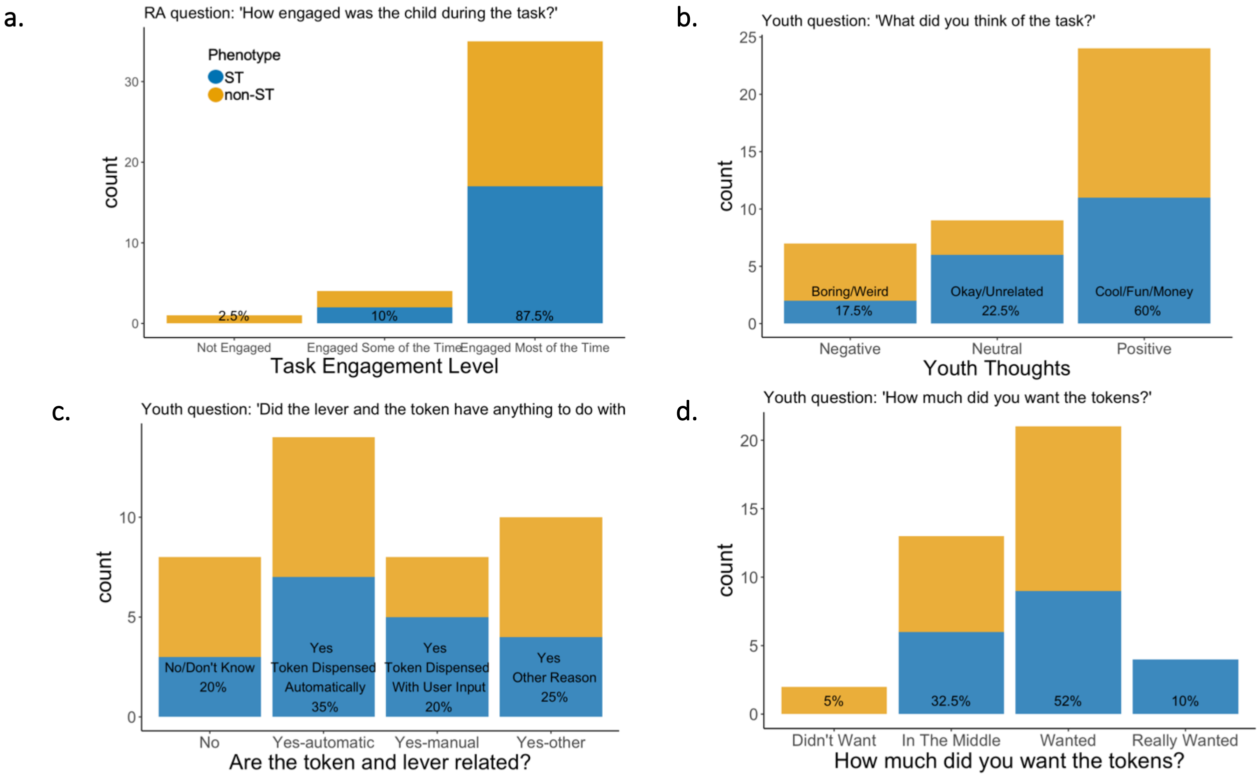


**Supplemental Figure S1. Split bar plot of Pavlovian conditioned approach task engagement.** Non-ST = non-sign-tracker, ST = sign-tracker. Percentages of the full sample are listed in each bar. Yellow split bars represent non-ST, blue split bars represent ST. **a.** research assistant (RA) reported post-session assessment of the level of engagement the child exhibited during the task. Answers were 'Not engaged', 'Engaged some of the time', or 'Engaged most of the time'. **b.** Youth-reported free-response question assessing what the participant thought of the task. Responses were categorized into negative responses (e.g., the task was boring or weird), neutral responses (e.g., the task was okay or made unrelated comments), or positive responses (e.g., the task was cool or fun, or they liked that they got money). **c.** youth-reported free-response question assessing whether the participant thought the token and lever were related. Responses were categorized into no (e.g., not related or don't know), yes-automatic (e.g., yes, the token dispensed automatically with no user input), yes-manual (yes, the token dispensed only with user input), and yes-other (yes they are related but the participant gave an unrelated answer, e.g., they are both green). **d.** youth-reported question assessing how much they wanted the tokens ranging from 'really didn't want', 'didn't want', 'in the middle', 'wanted', or 'really wanted'.

Supplemental Table S1. Lever- and reward-directed behavior group means and standard deviations by block

a. Conditioned stimulus period normalized by time of each trial

| | non-ST | | | | ST | | | |
|---|---|---|---|---|---|---|---|---|
| Block | 1 | 2 | 3 | 4 | 1 | 2 | 3 | 4 |
| Lever Contacts | 0.13 (0.23) | 0.11 (0.20) | 0.04 (0.08) | 0.05 (0.09) | 0.73 (0.76) | 1.02 (0.89) | 1.04 (0.73) | 0.96 (0.71) |
| Lever Probability | 0.31 (0.04) | 0.03 (0.04) | 0.02 (0.02) | 0.02 (0.03) | 0.09 (0.04) | 0.10 (0.03) | 0.11 (0.02) | 0.11 (0.02) |
| Lever Latency | 6.70 (2.29) | 6.64 (2.54) | 7.30 (1.87) | 7.15 (1.96) | 4.12 (2.63) | 3.41 (2.08) | 3.06 (1.92) | 2.96 (1.33) |
| Reward Contacts | 0.01 (0.02) | 0.01 (0.01) | 0.02 (0.06) | 0.01 (0.03) | 0.02 (0.03) | 0.05 (0.10) | 0.03 (0.07) | 0.02 (0.07) |
| Reward Probability | 0.00 (0.01) | 0.00 (0.01) | 0.01 (0.02) | 0.01 (0.01) | 0.01 (0.01) | 0.01 (0.01) | 0.00 (0.01) | 0.00 (0.00) |
| Reward Latency | 7.84 (0.93) | 7.63 (1.83) | 7.32 (1.91) | 7.47 (1.86) | 8.00 (0.51) | 7.70 (1.20) | 8.02 (0.45) | 8.06 (0.59) |
| Response Bias | 0.24 (0.29) | 0.19 (0.34) | 0.06 (0.17) | 0.09 (0.21) | 0.66 (0.40) | 0.77 (0.30) | 0.89 (0.15) | 0.88 (0.16) |
| PavCA Index | 0.20 (0.27) | 0.17 (0.31) | 0.04 (0.16) | 0.08 (0.18) | 0. 06 (0.38) | 0.68 (0.29) | 0.79 (0.16) | 0.79 (0.16) |

b. Intertrial interval period normalized by time of each trial

| | non-ST | | | | ST | | | |
|---|---|---|---|---|---|---|---|---|
| Block | 1 | 2 | 3 | 4 | 1 | 2 | 3 | 4 |
| Lever Contacts | 0.01 (0.02) | 0.03 (0.06) | 0.01 (0.03) | 0.01 (0.03) | 0.06 (0.14) | 0.02 (0.03) | 0.05 (0.07) | 0.04 (0.06) |
| Lever Probability | 0.00 (0.01) | 0.00 (0.01) | 0.00 (0.00) | 0.00 (0.01) | 0.01 (0.01) | 0.01 (0.01) | 0.01 (0.01) | 0.01 (0.01) |
| Lever Latency | 19.80 (3.64) | 18.30 (5.11) | 18.60 (5.04) | 19.80 (5.41) | 20.90 (2.85) | 20.70 (2.16) | 20.80 (2.53) | 20.30 (2.81) |
| Reward Contacts | 0.07 (0.05) | 0.07 (0.04) | 0.07 (0.03) | 0.06 (0.04) | 0.11 (0.09) | 0.11 (0.12) | 0.08 (0.05) | 0.08 (0.04) |
| Reward Probability | 0.04 (0.02) | 0.05 (0.03) | 0.04 (0.03) | 0.04 (0.02) | 0.04 (0.02) | 0.04 (0.02) | 0.04 (0.02) | 0.05 (0.02) |
| Reward Latency | 19.00 (4.14) | 17.80 (5.11) | 16.7 (5.86) | 18.3 (5.58) | 19.9 (3.62) | 18.9 (4.35) | 20.1 (2.22) | 19.5 (2.68) |
| Response Bias | -0.71 (0.38) | -0.63 (0.44) | -0.65 (0.37) | -0.67 (0.38) | -0.54 (0.46) | -0.61 (0.41) | -0.58 (0.42) | -0.67 (0.38) |
| PavCA Index | -0.48 (0.26) | -0.43 (0.30) | -0.46 (0.24) | -0.47 (0.25) | -0.38 (0.31) | -0.44 (0.29) | -0.40 (0.28) | -0.46 (0.25) |

c. Conditioned stimulus period non-normalized values

| Block | non-ST 1 | non-ST 2 | non-ST 3 | non-ST 4 | ST 1 | ST 2 | ST 3 | ST 4 |
|---|---|---|---|---|---|---|---|---|
| Lever Contacts | 1.07 (1.91) | 0.90 (1.70) | 0.35 (0.64) | 0.42 (0.72) | 6.06 (6.29) | 8.46 (7.35) | 8.60 (6.08) | 8.00 (5.85) |
| Lever Probability | 0.25 (0.30) | 0.22 (0.33) | 0.13 (0.18) | 0.16 (0.22) | 0.70 (0.35) | 0.81 (0.24) | 0.91 (0.12) | 0.90 (0.13) |
| Lever Latency | 6.70 (2.29) | 6.64 (2.54) | 7.30 (1.87) | 7.15 (1.96) | 4.12 (2.63) | 3.41 (2.08) | 3.06 (1.92) | 2.96 (1.33) |
| Reward Contacts | 0.06 (0.16) | 0.05 (0.11) | 0.19 (0.50) | 0.11 (0.23) | 0.16 (0.25) | 0.45 (0.87) | 0.24 (0.60) | 0.19 (0.59) |
| Reward Probability | 0.02 (0.05) | 0.03 (0.07) | 0.08 (0.14) | 0.07 (0.11) | 0.04 (0.09) | 0.04 (0.10) | 0.02 (0.04) | 0.02 (0.03) |
| Reward Latency | 7.84 (0.93) | 7.63 (1.83) | 7.32 (1.91) | 7.47 (1.86) | 8.00 (0.51) | 7.70 (1.20) | 8.02 (0.45) | 8.06 (0.59) |
| Response Bias | 0.24 (0.29) | 0.19 (0.34) | 0.06 (0.17) | 0.09 (0.21) | 0.66 (0.40) | 0.77 (0.30) | 0.89 (0.15) | 0.88 (0.16) |
| PavCA Index | 0.20 (0.27) | 0.17 (0.31) | 0.04 (0.16) | 0.08 (0.18) | 0.60 (0.38) | 0.68 (0.29) | 0.79 (0.16) | 0.79 (0.16) |

*Note.* means and standard deviations of reward- and lever-directed responses for non-STs and STs. non-ST = non-sign-tracker; ST = sign-tracker; PavCA = Pavlovian Conditioned Approach. **a.** values were normalized by dividing each score by the length the phase (8 seconds for conditioned stimulus). **b.** values were normalized by dividing each score by the length of each respective intertrial interval (8, 16, 24, or 32 seconds for each ITI). **c.** raw non-normalized values during the conditioned stimulus period.

Supplemental Table S2. Lever- and Reward-Directed Behavior Correlations (block 3/4 average)

| | CS | | | | | | ITI | | | | |
|---|---|---|---|---|---|---|---|---|---|---|---|
| | Lever Contacts | Reward Contacts | Lever Latency | Reward Latency | Lever Probability | Reward Probability | Lever Contacts | Reward Contacts | Lever Latency | Reward Latency | Lever Probability |
| CS | | | | | | | | | | | |
| Lever Contacts | | | | | | | | | | | |
| Reward Contacts | 0.23 | | | | | | | | | | |
| Lever Latency | -0.73*** | -0.13 | | | | | | | | | |
| Reward Latency | 0.18 | -0.08 | 0.22 | | | | | | | | |
| Lever Probability | 0.75*** | 0.16 | -0.83*** | 0.24 | | | | | | | |
| Reward Probability | -0.26 | 0.32* | 0.24 | -0.20 | -0.18 | | | | | | |
| ITI | | | | | | | | | | | |
| Lever Contacts | 0.59*** | 0.15 | -0.42** | 0.01 | 0.41** | 0.15 | | | | | |
| Reward Contacts | 0.03 | 0.07 | -0.08 | -0.01 | 0.24 | 0.34* | 0.17 | | | | |
| Lever Latency | 0.16 | 0.08 | 0.26 | 0.82*** | 0.21 | 0.02 | 0.08 | 0.33* | | | |
| Reward Latency | 0.21 | -0.08 | 0.15 | 0.91*** | 0.26 | -0.29 | -0.04 | 0.06 | 0.88*** | | |
| Lever Probability | 0.60*** | 0.20 | -0.44** | -0.01 | 0.45** | 0.21 | 0.92*** | 0.18 | 0.07 | -0.05 | |
| Reward Probability | -0.07 | -0.14 | 0.13 | 0.41** | 0.11 | -0.13 | -0.32* | 0.27 | 0.41** | 0.42** | -0.34* |

*Note.* CS = conditioned stimulus phase with lever presentation; ITI = inter-trial interval phase; $p < .001$***, $p < .01$**, $p < .05$*

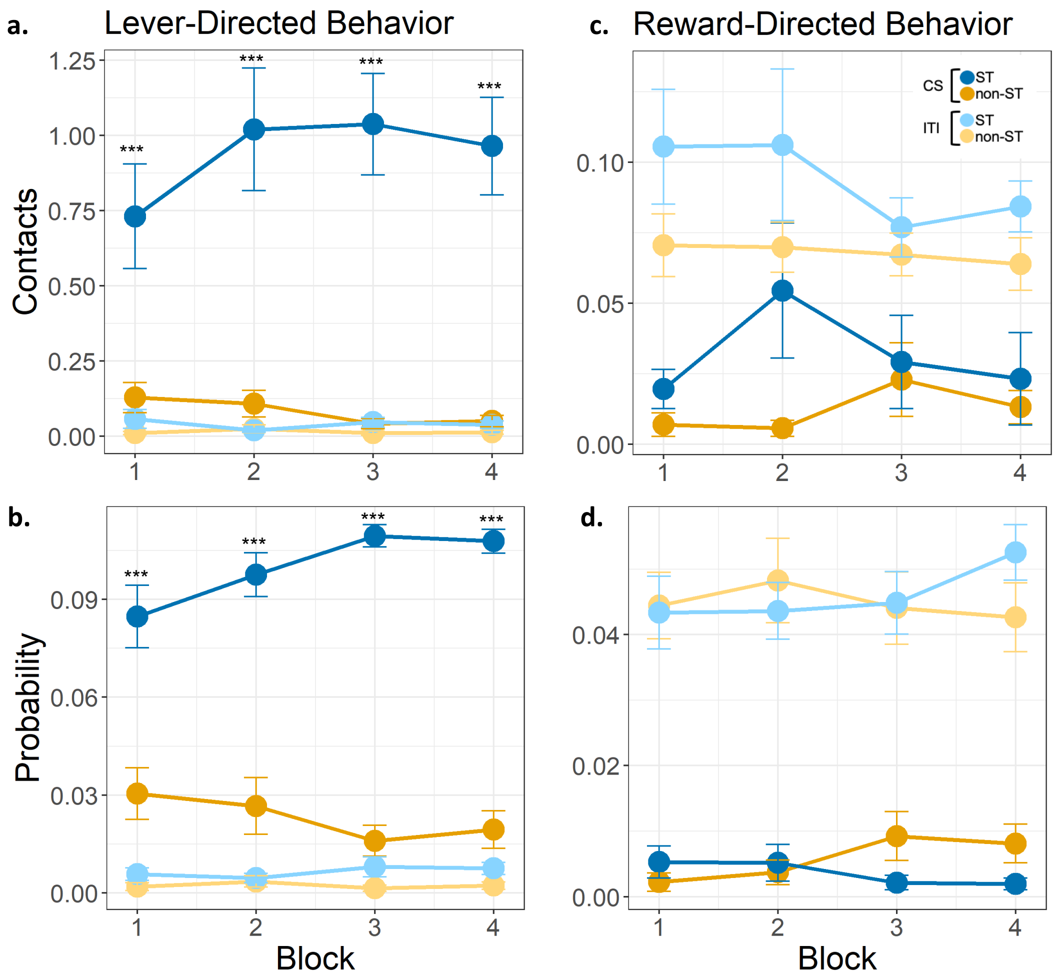


**Supplemental Figure S2. Plots for lever- and reward-directed behaviors normalized by time.** Behaviors across all 4 blocks between phenotypes (ST, non-ST) and between phases (CS, ITI); CS = conditioned stimulus/lever presentation; ITI = inter-trial interval; ST = sign-tracker; non-ST = non-sign-tracker. Values were normalized by dividing each score by the length the phase (8 seconds for conditioned stimulus and 8, 16, 24, or 32 seconds for each intertrial interval). Error bars represent standard error of the mean. Post-hoc tests between phenotypes during CS for each block are marked with asterisks; $p < .001$***, $p < .01$**, $p < .05$* **a.** Lever contacts; significant main effect for phase, $F_{1,266} = 193.34$, $p < .001$, and phenotype, $F_{1,36} = 29.08$, $p < .001$; significant interaction effect for phenotype by phase, $F_{1,266} = 142.82$, $p < .001$. **b.** Lever probability; significant main effect for sex, $F_{1,36} = 6.31$, $p = .015$, phase, $F_{1,266} = 738.13$, $p < .001$, and phenotype, $F_{1,36} = 106.47$, $p < .001$; significant interaction effect for phenotype by phase, $F_{1,266} = 296.82$, $p < .001$, phenotype by time, $F_{1,266} = 5.30$, $p = .001$, and phenotype by time by phase $F_{3,266} = 3.84$, $p = .01$. **c.** Reward contacts; significant main effect for age, $F_{1,36} = 4.33$, $p = .044$, and phase, $F_{1,266} = 103.60$, $p < .001$. **d.** Reward probability; significant main effect for age, $F_{1,36} = 6.62$, $p = .01$, phase, $F_{1,266} = 504.78$, $p < .001$, and sex, $F_{1,36} = 7.43$, $p = .009$.

Supplemental Table S3. Symptom and environmental differences by phenotype

| | non-ST (N=21) | ST (N=19) | *t*-statistic | *p*-value | Cohen's d | df | se | conf from | conf to |
|---|---|---|---|---|---|---|---|---|---|
| Age in Months | 120 (12.5) | 119 (12.2) | 0.22 | 0.824 | 0.07 | 37.795 | 3.913 | -7.046 | 8.8 |
| Social Problems (CBCL) | -0.479 (0.782) | 0.459 (1.00) | -3.1 | 0.004 | 1.05 | 30.164 | 0.302 | -1.556 | -0.321 |
| Thought Problems (CBCL) | -0.438 (0.807) | 0.264 (1.08) | -2.2 | 0.036 | 0.75 | 29.52 | 0.32 | -1.356 | -0.049 |
| Somatic Complaints (CBCL) | -0.156 (0.917) | -0.0596 (1.07) | -0.29 | 0.775 | 0.1 | 31.803 | 0.334 | -0.776 | 0.583 |
| Attention Problems (CBCL) | -0.430 (0.946) | 0.403 (0.921) | -2.68 | 0.011 | 0.89 | 33.739 | 0.312 | -1.467 | -0.201 |
| Rule Breaking Behavior (CBCL) | -0.348 (0.863) | 0.153 (1.03) | -1.57 | 0.126 | 0.53 | 31.366 | 0.319 | -1.152 | 0.149 |
| Aggressive Behavior (CBCL) | -0.316 (0.722) | 0.484 (1.17) | -2.44 | 0.022 | 0.84 | 26.126 | 0.328 | -1.473 | -0.126 |
| Oppositional Defiance (CBCL-DSM) | -0.470 (0.770) | 0.523 (0.943) | -3.44 | 0.002 | 1.16 | 30.981 | 0.289 | -1.582 | -0.404 |
| Attention Deficit Hyperactivity (CBCL-DSM) | -0.419 (0.819) | 0.401 (1.01) | -2.66 | 0.012 | 0.9 | 30.94 | 0.308 | -1.447 | -0.191 |
| Anxious Depressed (CBCL-DSM) | 0.0595 (1.09) | -0.113 (0.996) | 0.5 | 0.623 | 0.16 | 33.986 | 0.348 | -0.535 | 0.881 |
| Withdrawn Depressed (CBCL-DSM) | -0.244 (0.984) | 0.101 (0.976) | -1.05 | 0.299 | 0.35 | 33.621 | 0.327 | -1.01 | 0.32 |
| Conduct Problems (CBCL-DSM) | -0.235 (0.807) | 0.091 (1.22) | -0.93 | 0.359 | 0.32 | 27.236 | 0.349 | -1.043 | 0.391 |
| Anxiety Problems (CBCL) | 0.121 (1.05) | -0.009 (1.08) | 0.37 | 0.716 | 0.12 | 33.382 | 0.355 | -0.592 | 0.853 |
| Depressive Problems (CBCL) | -0.287 (0.969) | 0.409 (0.848) | -2.3 | 0.028 | 0.76 | 33.988 | 0.303 | -1.311 | -0.081 |
| Somatic Problems (CBCL) | -0.226 (0.872) | -0.034 (1.01) | -0.61 | 0.546 | 0.21 | 31.902 | 0.316 | -0.836 | 0.451 |
| Internalizing Problems (CBCL) | 48.1 (10.9) | 48.5 (9.25) | -0.11 | 0.914 | 0.04 | 33.927 | 3.355 | -7.185 | 6.454 |
| Externalizing Problems (CBCL) | 43.3 (7.53) | 50.8 (9.33) | -2.64 | 0.013 | 0.89 | 30.786 | 2.846 | -13.32 | -1.701 |
| Activation Control (EATQ) | 3.25 (0.823) | 3.07 (0.616) | 0.78 | 0.44 | 0.25 | 36.402 | 0.231 | -0.288 | 0.649 |
| Affiliation (EATQ) | 4.10 (0.385) | 3.96 (0.604) | 0.85 | 0.404 | 0.28 | 28.041 | 0.165 | -0.198 | 0.479 |

| | non-ST (N=21) | ST (N=19) | *t*-statistic | *p*-value | Cohen's d | df | se | conf from | conf to |
|---|---|---|---|---|---|---|---|---|---|
| Aggressive Behaviors (EATQ) | 2.02 (0.548) | 2.67 (0.740) | -3.06 | 0.005 | 1 | 30.942 | 0.211 | -1.078 | -0.215 |
| Attention (EATQ) | 3.46 (0.715) | 3.07 (0.655) | 1.76 | 0.087 | 0.56 | 36.815 | 0.219 | -0.058 | 0.831 |
| Depressive Behaviors (EATQ) | 1.89 (0.463) | 2.49 (0.599) | -3.47 | 0.001 | 1.14 | 31.784 | 0.174 | -0.957 | -0.25 |
| Fear (EATQ) | 2.57 (0.544) | 3.08 (0.592) | -2.79 | 0.008 | 0.9 | 34.953 | 0.183 | -0.884 | -0.14 |
| Frustration (EATQ) | 2.62 (0.388) | 3.03 (0.667) | -2.29 | 0.03 | 0.76 | 26.388 | 0.179 | -0.776 | -0.042 |
| Inhibition (EATQ) | 3.89 (0.564) | 3.56 (0.379) | 2.17 | 0.037 | 0.68 | 35.151 | 0.152 | 0.021 | 0.639 |
| Shyness (EATQ) | 2.50 (0.920) | 2.79 (0.827) | -1.02 | 0.317 | 0.32 | 36.899 | 0.28 | -0.851 | 0.283 |
| Surgency (EATQ) | 3.44 (0.750) | 3.28 (0.462) | 0.82 | 0.42 | 0.25 | 33.836 | 0.197 | -0.239 | 0.56 |
| Negative Affect (EATQ) | 2.18 (0.367) | 2.73 (0.598) | -3.41 | 0.002 | 1.14 | 27.318 | 0.162 | -0.885 | -0.22 |
| Surgency Composite (EATQ) | 3.46 (0.592) | 3.14 (0.470) | 1.87 | 0.069 | 0.59 | 36.816 | 0.17 | -0.026 | 0.664 |
| Perseverance (UPPS-P) | 7.48 (2.27) | 7.39 (2.75) | 0.11 | 0.915 | 0.03 | 33.106 | 0.815 | -1.572 | 1.746 |
| Positive Urgency (UPPS-P) | 9.48 (2.32) | 8.61 (3.20) | 0.95 | 0.348 | 0.31 | 30.456 | 0.908 | -0.989 | 2.719 |
| Negative Urgency (UPPS-P) | 10.3 (2.33) | 9.94 (2.69) | 0.48 | 0.635 | 0.16 | 33.97 | 0.813 | -1.263 | 2.041 |
| Sensation Seeking (UPPS-P) | 10.3 (2.28) | 9.83 (2.26) | 0.62 | 0.539 | 0.2 | 36.225 | 0.729 | -1.025 | 1.93 |
| Planning (UPPS-P) | 7.86 (1.77) | 8.06 (2.34) | -0.29 | 0.77 | 0.1 | 31.355 | 0.673 | -1.57 | 1.173 |
| Behavioral Activation: Drive | 3.48 (2.06) | 5.11 (3.31) | -1.82 | 0.08 | 0.6 | 27.64 | 0.9 | -3.48 | 0.21 |
| Behavioral Activation: Fun | 6.19 (2.25) | 5.67 (2.77) | 0.64 | 0.525 | 0.21 | 32.791 | 0.816 | -1.137 | 2.184 |
| Behavioral Activation: Reward | 11.2 (2.34) | 11.1 (3.08) | 0.14 | 0.887 | 0.05 | 31.437 | 0.889 | -1.685 | 1.939 |
| Behavioral Inhibition (BIS/BAS) | 11.2 (2.34) | 11.5 (3.79) | -0.3 | 0.766 | 0.1 | 27.401 | 1.029 | -2.42 | 1.801 |
| Crystalized Cognitive Functioning (NIH toolbox) | 48.1 (10.8) | 43.5 (9.88) | 1.36 | 0.183 | 0.44 | 35.982 | 3.35 | -2.245 | 11.345 |

| | non-ST (N=21) | ST (N=19) | *t*-statistic | *p*-value | Cohen's d | df | se | conf from | conf to |
|---|---|---|---|---|---|---|---|---|---|
| Fluid Cognitive Functioning (NIH toolbox) | 46.1 (11.3) | 42.8 (10.5) | 0.91 | 0.367 | 0.3 | 34.646 | 3.585 | -4.005 | 10.558 |
| Household Income | 4.55 (0.759) | 3.44 (1.50) | 2.81 | 0.01 | 0.94 | 24.538 | 0.393 | 0.295 | 1.916 |
| PACES | 8.67 (1.35) | 7.53 (1.42) | 2.51 | 0.017 | 0.82 | 33.648 | 0.454 | 0.215 | 2.06 |

*Note.* CBCL = Child Behavior Checklist (parent report); EATQ = Early Adolescent Temperament Questionnaire (parent report); UPPS-P = Urgency, Premeditation, Perseverance, Sensation Seeking, Positive Urgency scale (youth self-report); BIS/BAS = Behavioral Inhibition/Behavioral Activation Scale (youth self-report); PACES = Protective and Compensatory Experiences Scale (parent report). Measures were tested for normality and all CBCL subscales (excluding Externalizing/Internalizing Problems scales) were log transformed. To directly compare symptoms between STs and non-STs, we used two-sided Welch two sample *t*-tests.

Supplemental Table S4. Pearson *r* Correlations among Symptoms and PavCA Scores

| | PavCA | Attention Deficit/ Hyperactivity Problems | Social Problems | Oppositional Defiance Problems | Fear Behaviors | Inhibitory Control |
|---|---|---|---|---|---|---|
| PavCA | | | | | | |
| Attention Deficit/Hyperactivity Problems (CBCL) | 0.34* | | | | | |
| Social Problems (CBCL) | 0.38* | 0.54*** | | | | |
| Oppositional Defiance Problems (CBCL) | 0.40* | 0.43** | 0.64*** | | | |
| Fear Behaviors (EATQ) | 0.39* | 0.23 | 0.26 | 0.48** | | |
| Inhibitory Control (EATQ) | -0.18 | -0.49** | -0.47** | -0.55*** | -0.09 | |
| Negative Affect (EATQ) | 0.42** | 0.29^ | 0.65*** | 0.80*** | 0.42** | -0.53*** |

*Note.* CBCL = Child Behavior Checklist; EATQ = Early Adolescent Temperament Questionnaire; PavCA = Pavlovian conditioned approach index; $p<.001$***, $p<.01$**, $p<.05$*, $p<.1$^

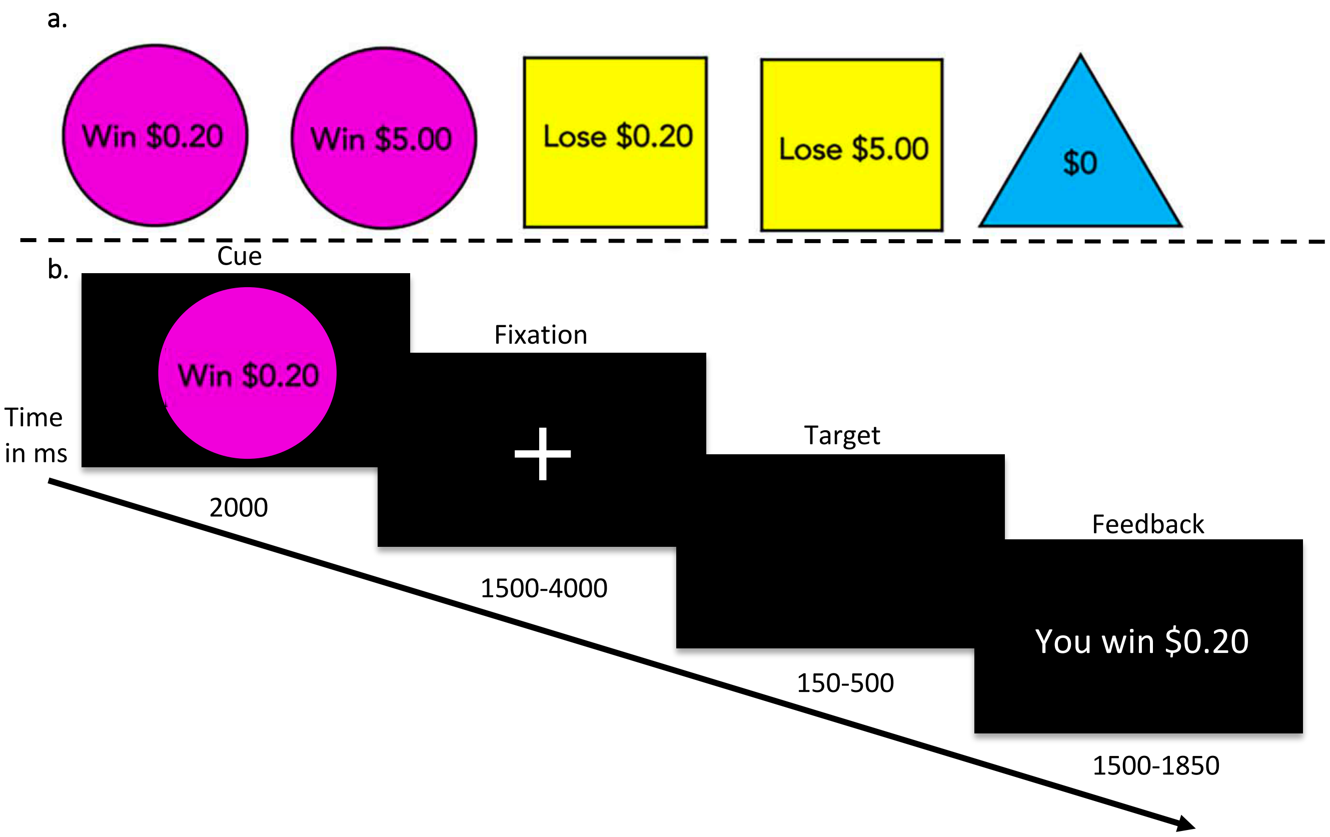


**Figure S3. Monetary incentive delay (MID) fMRI task cues and timeseries. a.** Stimuli used in the monetary incentive delay (MID) task included small win (win $0.20), large win (win $5.00), small loss (lose $0.20), large loss (lose $5.00) or neutral (no win or loss). **b.** Timeseries of events. Cues signaling task condition were displayed for 2 seconds followed by a 1.5 to 4 second fixation cross and then a black target screen for .15 to .5 seconds. Participants were instructed to press a button while the target was on the screen. Participants would neither gain nor lose money for missing the target. The outcome of each trial was presented immediately after each response for 2 seconds minus the duration of the target.

Supplemental Table S5. Regions with group by condition differences in voxel-wise whole brain analysis

| MID Contrast | Region | # Voxels | CM x | CM y | CM z | Pattern |
|---|---|---|---|---|---|---|
| win-neutral | Left Inferior Parietal Lobe | 353 | 27.3 | 56.3 | 42.2 |  |
| win-neutral | Left Inferior Parietal Lobe | 138 | 43.0 | 59.7 | 14.0 | 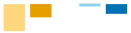 |
| win-neutral | Right Inferior Parietal Lobe | 59 | -29.5 | 44.1 | 36.6 | 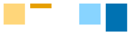 |
| win-neutral | Right Middle Frontal Gyrus | 45 | -22.8 | -0.3 | 42.7 | 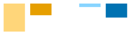 |
| win-neutral | Right Amygdala | 60 | -24.6 | 3.2 | -13.7 | 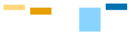 |
| win-neutral | Left Pre/Postcentral Gyrus | 56 | 20.7 | 25.6 | 53.1 | 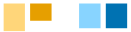 |
| loss-neutral | Left Inferior/Superior Parietal Lobe | 120 | 29.0 | 46.7 | 42.0 | 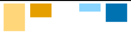 |
| loss-neutral | Left Posterior Cingulate | 64 | 1.2 | 13.3 | 39.6 | 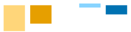 |
| loss-neutral | Left Precentral Gyrus | 40 | 46.9 | 5.0 | 43.1 | 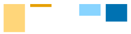 |
| loss-neutral | Left Pre/Postcentral Gyrus | 61 | 22.9 | 25.5 | 55.5 | 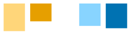 |
| loss-neutral | Right Superior Frontal Gyrus | 148 | -0.3 | -7.2 | 51.5 | 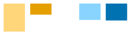 |
| loss-neutral | Right Superior Frontal Gyrus | 42 | -0.2 | -48.9 | 19.4 | 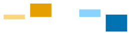 |
| loss-neutral | Right Anterior Cingulate/Superior Frontal | 84 | -2.1 | -34.3 | 23.1 | 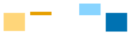 |
| loss-neutral | Right Medial Orbital Frontal Gyrus | 62 | -3.2 | -47.9 | 2.5 | 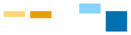 |
| loss-neutral | Left Bankssts | 46 | 54.3 | 55.1 | 15.7 | 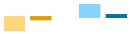 |

*Note.* All results corrected for multiple comparisons at $p < 0.05$. Significant clusters with group by condition effects in voxel-wise whole brain analysis in the high-win versus neutral and high-loss versus neutral conditions. Number of voxels and center mass coordinates from the Haskins Pediatric atlas are listed for each cluster. The pattern includes a bar graph representing percent blood oxygen level dependent (BOLD) signal change during each condition by phenotype. Yellow represents non-sign-trackers (non-ST) and blue represents sign-trackers (ST) with lighter shades indicating neutral trials and darker shades indicating win/loss trials respectively (e.g., non-ST neutral, non-ST win, ST neutral, ST win). Bar graphs in the *pattern* column reflect those in Figure 3.

Supplemental Table S6: Correlations between Brain Activations and Behaviors

| | Win-neutral | | Loss-neutral |
|---|---|---|---|
| | Left IPL | Right Amygdala | Left IPL |
| Win-neutral Left IPL | | | |
| Win-neutral Right Amygdala | -0.28 | | |
| Loss-neutral Left IPL | 0.71*** | -0.14 | |
| Oppositional Defiant Problems (CBCL) | -0.43* | 0.00 | -0.20 |
| Externalizing Problems (CBCL) | -0.36^ | 0.00 | -0.17 |
| Inhibitory Control (EATQ) | 0.36^ | -0.17 | 0.15 |
| Surgency (EATQ) | -0.28 | -0.27 | -0.45* |
| Negative Affect (EATQ) | -0.34^ | 0.05 | -0.21 |
| Sensation Seeking (UPPS-P) | -0.33^ | 0.32^ | -0.23 |
| PavCA Index | -0.46* | 0.48** | -0.45* |

*Note.* IPL = inferior parietal lobe; CBCL = Child Behavior Checklist; UPPS-P = Urgency, Premeditation, Perseverance, Sensation Seeking, Positive Urgency scale; EATQ = Early Adolescent Temperament Questionnaire; PavCA Index = Pavlovian conditioned approach index; p<.001***, p<.01**, p<.05*, p<.1^

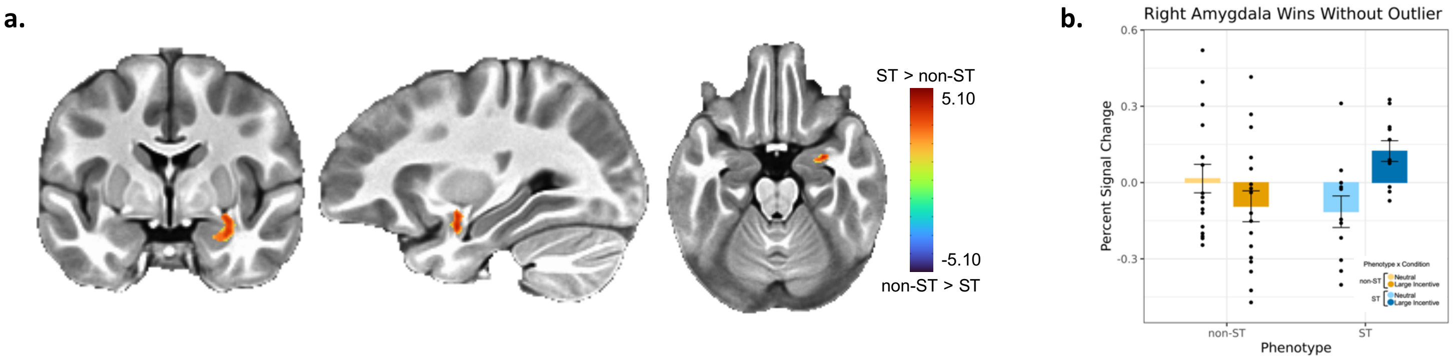


**Supplemental Figure S4: Sensitivity analysis for amygdala activation for functional magnetic resonance imaging (fMRI) percent BOLD signal change during the Monetary Incentive Delay (MID) task for win anticipation.** ST = sign-tracker (n = 11), non-ST = non-sign-tracker (n = 17). Results are based on whole-brain voxelwise linear mixed effects with fixed effects for group, condition, age, sex, a group by condition interaction and a random intercept for subject. We followed this with planned contrasts investigating a group (ST, non-ST) by condition (win-neutral) interaction. As a supplementary sensitivity analysis, one potential outlier in the ST group was removed. The effect remains consistent with the full sample results reported in the main text but is visible at a less conservative threshold with more prominent post hoc group differences during the win condition ($t_{45}$ = -2.89, p = .029). **a.** Significant clusters reported using a voxelwise *p*-value threshold of 0.007 and α < 0.05 at the cluster level (N = 11.73 voxels). Estimated marginal means were used for post-hoc tests. Brain activation colors represent *t*-test *z*-statistics for group differences in percent signal change between conditions (win-neutral). Blue represents negative values and indicates a greater percent signal change for non-ST than ST between neutral and large win condition. Red represents positive values that indicate greater percent signal change for STs between neutral and large win condition. **b.** Bar graph depicts percent signal change in respective clusters across neutral and large win condition. Non-STs are represented by yellow bars and STs by blue. Lighter colors indicate neutral trials whereas darker colors indicate win trials. Significant differences derived from post hoc tests are marked with asterisks; *p* < .01**, *p* < .05*. Significant contrasts (win-neutral) correspond with brain images such that blue indicates a significant contrast for non-ST and red indicates a significant contrast for STs. BOLD activations for a group by condition interaction in the right amygdala during win anticipation compared to neutral anticipation ($F_{1,26}$ = 12.31, *p* = .002, $\eta^2_p$ = 0.32, 90% CI = 0.09–0.52). Post hoc tests indicate that, in contrast to the IPL, STs significantly increase BOLD activation from neutral to win conditions ($t_{26}$ = -3.09, *p* = .023, Cohen's d = -1.32, s.e. = 0.45, 90% CI = -2.24 – -0.40) indicating salience-dependent modulation within the amygdala. Again, in contrast to the IPL, non-STs did not differ between conditions or from STs during the neutral condition (*p*s > .03). They did differ from STs during the win condition ($t_{45}$ = -2.89, *p* = .029, Cohen's d = -1.33, s.e. = 0.48, 90% CI = -2.30 – -0.36).
